# Mitf links neuronal activity and long-term homeostatic intrinsic plasticity

**DOI:** 10.1101/507640

**Authors:** Diahann A. M. Atacho, Hallur Reynisson, Anna Þóra Pétursdóttir, Thor Eysteinsson, Eiríkur Steingrímsson, Pétur Henry Petersen

## Abstract

Neuroplasticity forms the basis for neuronal circuit complexity and differences between otherwise similar circuits. We show that the Microphthalmia-associated transcription factor (*Mitf*) plays a central role in intrinsic plasticity of olfactory bulb (OB) projection neurons. Mitral and tufted (M/T) neurons from *Mitf* mutant mice are hyperexcitable, have a reduced Type-A potassium current (I_A_) and exhibit reduced expression of *Kcnd3*, which encodes a potassium voltage-gated channel subunit (Kv4.3) important for generating the I_A_. Furthermore, expression of the *Mitf* and *Kcnd3* genes is activity-dependent in OB projection neurons, The MITF protein activates expression from *Kcnd3* regulatory elements. Moreover, *Mitf* mutant mice have changes in olfactory habituation and have increased habitutation for an odourant following long-term exposure, indicating that regulation of *Kcnd3* is pivotal for long-term olfactory adaptation. Our findings show that *Mitf* acts as a direct regulator of intrinsic homeostatic feedback and links neuronal activity, transcriptional changes and neuronal function.

**Significance statement:** These findings broaden the general understanding of transcriptional regulation of intrinsic plasticity in learning and memory. Regulation of intrinsic plasticity has wide-ranging implications and fundamental importance for neurological diseases.

## Introduction

Neuronal plasticity is comprised of activity-dependent changes that alter synaptic or intrinsic excitability of neurons [1], affecting the neuronal response, the gain control relationship between input and output. Ultimately it connects neuronal activity with changes in neuronal behavior [1, 2]. It can also be homeostatic, i.e. neurons with increased activity becoming less sensitive and neurons with lowered activity more sensitive [3–6]. Synaptic plasticity and intrinsic plasticity are thought to work at different time-scales and to interact and complement each other [7, 8]. Although synaptic plasticity is fairly well characterized, much less is known about the molecular mechanisms underlying intrinsic plasticity and its transcriptional regulation [9, 10].

The olfactory system is an attractive system to study plasticity, due to its clear flow of information, well-defined neuronal subtypes, ease of activation with odourants and well defined neuronal circuits. The OB is the first relay of peripheral olfactory information within the CNS. Each olfactory bulb in the mouse contains around 2000 glomeruli, spherical neuropil structures connecting peripheral olfactory sensory input and cortical structures. In the OB, two types of projection neurons are designated to each glomerulus. They are activated by olfactory sensory neurons (OSN) and target other CNS regions. These are the mitral cells (MC) [11, 12] located in the mitral cell layer (MCL) and the tufted neurons located in the external plexiform layer (EPL). These neurons show remarkable capacity in normalizing sensory input, by amplifying low signals and reducing high signals [13]. Odour exposure also reduces the sensitivity of mitral and tufted cells (M/T) [14, 15]. In addition, it has been shown that there is high inter-glomerular functional diversity [16], showing that each glomerulus is, to a large degree, regulated independently by its activity. The OB therefore shows clear changes in gain-control and homeostasic plasticity and it has been suggested that intrinsic changes are fundamental to OB sensory adaptation [17].

The transcription factor MITF, best known as a master regulator of melanocytes [18], is specifically expressed in the M/T neurons of the OB [19]. As the M/T neurons are the key output neurons in the OB circuitry, *Mitf* might play a role in olfaction [19]. Mitf is also expressed in a subset of tufted cells, the external tufted cells (ETC), which are excitatory and synapse with MC, but do not project out of the OB [20–22]. Here we show that Mitf links neuronal activity and intrinsic activity-dependent changes in the OB by regulating the expression of the potassium channel subunit *Kcnd3*, leading to a homeostatic response in OB’s projection neurons. Therefore, we suggest that *Mitf* plays a key role in olfactory adaptation and intrinsic homeostatic plasticity.

## Results

### *Mit* mutant mice have increased numbers of excititatory neurons

In order to characterize *Mitf* expression in the OB, we used multiplex fluorescent RNA *in situ* hybridization (mFISH) which generates a single fluorescent dot per transcript detected [23], allowing for single-cell analysis of gene expression. Consistent with previous observations [19], this showed that in wild type mice *Mitf* is expressed in M/T neurons, including in external tufted cells (ETC) located in the external plexiform layer (Figure 1a, b) and at low levels in the granule cell layer (GCL) (Figure 1a, arrowheads). Interestingly, there is significantly more *Mitf* expressed in the ETCs compared to MCs (F(1,15)=8.94, p=0.0092, Two-way ANOVA). Mice homozygous for the *Mitf^mi-vga9^* mutation are white and microphthalmic due to a transgene insertion mutation that affects expression of *Mitf* [24]. As expected, *Mitf* expression was decreased in the ETCs (t(14)=7.923, p<0.0001, Sidak multiple comparison test) and MCs (t(14)=2.969, p=0.0202, Sidak multiple comparison) in OBs of *Mitf* mutant mice (Figure 1 b,c).

**Figure 1:**
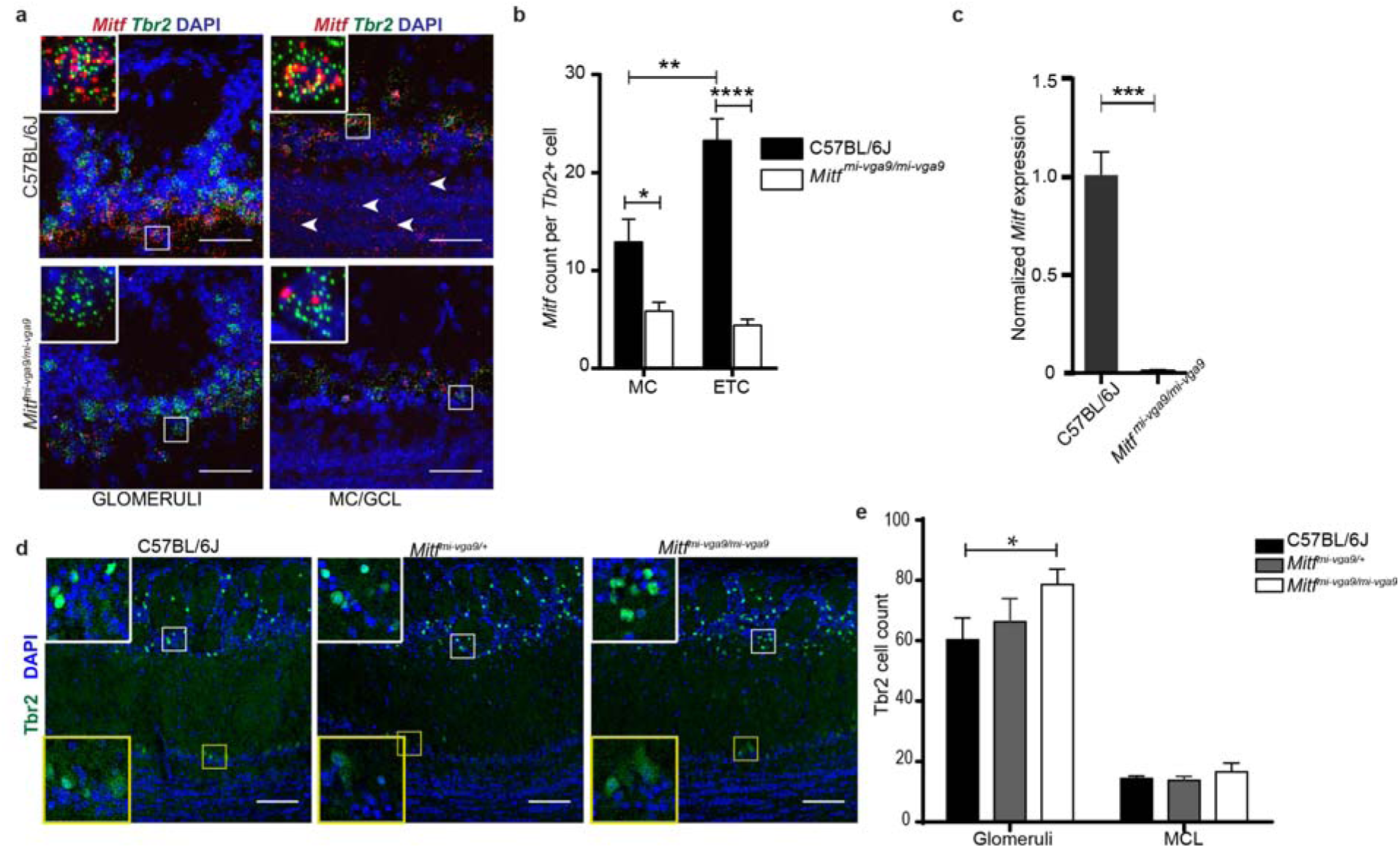
*Mitf* mutant mice have an increase in Tbr+ neurons in the glomerular layer. **a.** RNA *in situ* hybridization of *Mitf* (red) and *Tbr2* (green) in glomeruli and MC/GCL of wild-type and *Mitf^mi-vga9/mi-vga9^* mice. Scale bars are 50µm. **b.** *Mitf* count per *Tbr2*-positive cells below glomeruli and MCL. N=9 per genotype. **c**. *Mitf* mRNA expression, determined by RT-qPCR. N=6 per genotype. **d.** Immunofluorescent staining of Tbr2/Eomes (Tbr2+) neurons (green) in the OBs of wild-type, *Mitf^mi-vga9/+^* and *Mitf^mi-vga9/mi-vga9^*mice. **e**. Cell count of Tbr2/Eomes (Tbr2+) cells in the glomeruli and MCL of wild-type, *Mitf^mi-vga9/+^* and *Mitf^mi-vga9/mi-vga9^* mice. N=6 per genotype. The values on the graphs are mean ± SEM. DAPI nuclear staining is shown in blue. Scale bars are 100µm. P-values were calculated using two-way ANOVA (**b,e**) or two-tailed unpaired student’s t-test (**c**). *= P<0.05, ***P<0.001. MC= mitral cells, MCL=mitral cell layer, EPL= external plexiform layer.

Histological analysis showed no detectable defects in the cellular architecture of OBs from P75-P95 *Mitf* mutant mice (Suppl. Figure 7). Expression of tyrosine hydroxylase (TH) is reduced upon loss of activity in the OB [25] but was not affected in the OBs of *Mitf* mutant mice (Suppl. Figure 7 b, c). This also suggests normal OB function. Importantly, *Tbr2*, a marker for M/T cells [26, 27], is expressed in *Mitf* mutant OBs (Figure 1a) showing normal OB ontogeny. Interestingly, cell counts showed an increase of glomerular Tbr2+ neurons (t(20)=2.914, p=0.0171, Sidak multiple comparison) in the glomeruli of *Mitf* mutant mice (Figure 1d,e). An increase in M/T neurons has been shown to occur with increased activity [28, 29]. Additionally, analysis of apoptosis showed a twofold increase in cell death in periglomerular cells of *Mitf* mutant mice (Suppl. Figure 8; t(16)=2.724, p=0.015, Two-tailed *t*-test) suggesting increased glomerular neuronal turnover in the mutant mice.

### *Mitf* mutant mice have reduced I_A_ and a concomittant increase in mitral cell activity

To examine whether the loss of *Mitf* resulted in altered M/T neuronal function, we performed whole-cell patch clamp analysis on cultured primary M/T neurons from wild-type and *Mitf* mutant mice, both under voltage clamp and current clamp conditions, bypassing possible secondary effects due to cortical feedback mechanisms or other compensatory mechanisms within the OB. To establish the identity of the cells in culture, several biophysical properties were determined (see methods). The likelihood of an action potential is decided by the balance between sodium influx and potassium efflux. The initial potassium efflux activated near the spiking threshold is termed the A-type potassium current (I_A_) and is a major determinant of the likelihood of an action potential. Patch clamp recordings under voltage clamp showed a reduction in the I_A_ K^+^-current in M/T neurons from *Mitf* mutant mice. (Figure 2a-d; t(50)=2.313, p=0.0491, Sidak multiple comparison). No significant reduction was observed in the I_DR_ current (Figure 2d). There was no significant correlation between the membrane resistance of the M/T neurons and the maximum recorded I_A_ K^+^-current or the I_DR_ current from either wild type or *Mitf* mutant mice. The I/V relationship denotes the relationship between the I_A_ K^+^-current density (pA/pF) across the membrane and the membrane potential, and thus takes size of the cells into account based on their measured membrane capacitance. The I/V curves show that the mutant neurons have an altered voltage dependence of activation of the I_A_ currents evoked (Figure 2e), with multiple comparisons between the means revealing that the current density is significantly reduced at depolarized membrane potentials, at −5 mV and higher (p>0.05). However, the channel kinetics, i.e. activation and inactivation of the channels mediating the isolated I_A_, remained the same in cells from wild-type and mutant mice (Suppl. Figure 9). Reduction in the I_A_ leads to increased likelihood of action potentials and hyperexcitability [30–32]. Accordingly, increased spiking was observed in the mutant neurons (Figure 2f), under current clamp configuration of the whole-cell patch clamp. The mean firing frequency of spikes in wild-type M/T neurons was 0.79±0.17 Hz, while it was 1.75±0.15 Hz in mutant M/T neurons (t(11)=1.61, p=0.007, Two-tailed *t*-test). The mean amplitude of action potentials was similar, or 62±4 mV in wild-type M/T neurons and 54±2.6 mV in mutant M/T neurons (t(11)=1.06, p=0.31, Two-tailed *t*-test). In addition to spikes, miniature excitatory postsynaptic potentials (EPSP) could be observed in the current clamp recordings. The mean frequency of miniature EPSPs in the wild-type M/T neurons was 0.27±0.04 Hz and 0.47±0.21 Hz (t(11)=−1.55, p=0.148, Two-tailed *t*-test) in mutant M/T neurons. Additionally, expression of *c-Fos*, an activity-dependent gene frequently used to monitor neuronal acivity, was increased in the *Mitf* mutant MCs (Fig. 2g, h; t(9)=3.093, p=0.0386, Sidak multiple comparison), indicating increased neuronal activity *in vivo*. A second, unidentified cell population in the GCL also showed clear increase in expression of *c-Fos* in the mutant (Fig. 2g, arrowheads).

**Figure 2:**
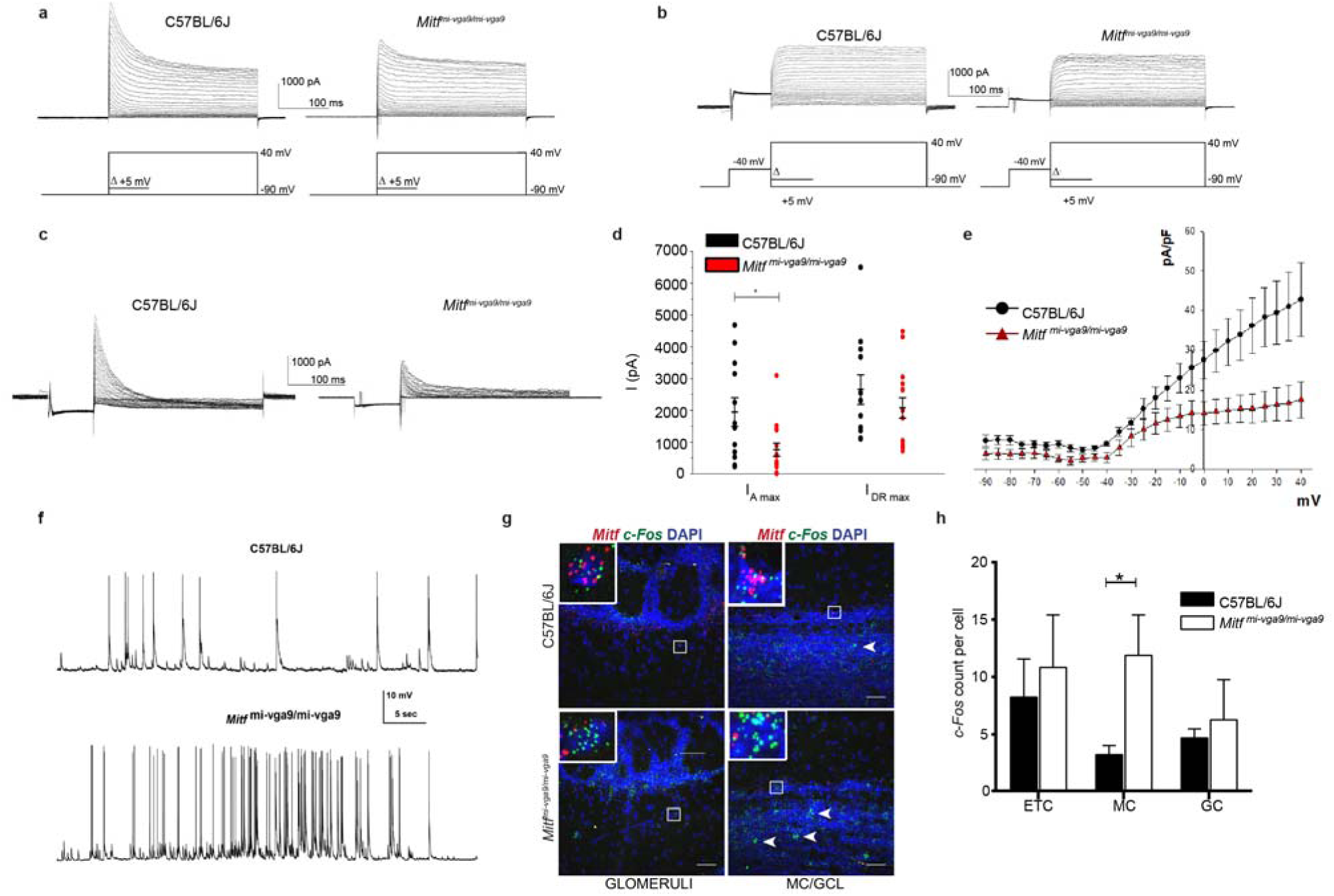
*Mitf* mutant mice have a decrease in I_A_ and a concomittant increase in mitral cell activity. **a**. Representative images of voltage clamp recordings. **b**. Representative images of voltage clamp recordings, where the I_A_ was inactivated by a −40mV pre-pulse. **c**. The isolated I_A_ recorded from wild-type and *Mitf^mi-vga9/mi-vga9^*M/T neurons. **d**. I_A max_ and I_DR max_ in wild-type and *Mitf^mi-vga9/mi-vga9^*. N=12 per genotype. **e**. The relation between current density (pA/pF) of isolated I_A_ currents and membrane voltage (mV) of wild-type and *Mitf^mi-vga9/mi-vga9^*M/T neurons. N=12 per genotype. **f**. Representative recordings of spontaneous action potentials observed in wild-type (N=5) and *Mitf^mi-vga9/mi-vga9^*M/T neurons (N=3) by current clamp. **g.** RNA *in situ* hybridization of *c-Fos* (green) and *Mitf* (red) in wild-type and *Mitf^mi-vga9/mi-vga9^* OBs. **h**. *c-Fos* dots per cell in the ETC, MC and GC. N=4 per genotype. The values on the graphs are mean ± SEM. DAPI nuclear staining is shown in blue. Scale bars are 50µm. P-values were calculated using two-tailed unpaired student’s t-test (**d**), non-linear regression and one-way ANOVA (**f**) and two-way ANOVA (**h**) and. *P<0.05. ETC= external tufted cells, MC= mitral cells, GC= granule cells.

### Expression of *Kcnd3* and *Mitf* is activity-dependent in the OB

Increased neuronal activity due to the loss of I_A_ led us to examine the expression of key potassium channel subunits in the OB. As displayed in Supplementary Table 1, many potassium channel subunits [33] are known to be expressed in the OB [34–36]. We focused our analysis on the I_A_-current regulating Kv4.3/*Kcnd3* potassium channel subunit, as according to ChIPseq data, MITF binds to its regulatory region (Supplementary Table 1). RNA *in situ* analysis showed reduced expression of *Kcnd3* in ETCs (t(12)=3.297, p=0.0190, Sidak multiple comparison) and MCs (t(12)=3.249, p=0.0208, Sidak multiple comparison) of *Mitf* mutant OBs, whereas the levels remained similar in cells of the GCL (Fig. 3a, b). Thus, the effects of *Mitf* on *Kcnd3* expression are cell autonomous, consistent with the possibility of *Mitf* regulating *Kcnd3* in M/T neurons. Although the ChIPseq data does not show MITF to bind to the regulatory region of the gene encoding the I_A_-regulating potassium channel subunit *Kcnd2* (Supplementary Table 1), its expression was increased in *Tbr2+* cells in OB from the *Mitf* mutant mouse (t(6)=5.051, p=0.0047, Sidak multiple comparison), possibly as a compensatory mechanism for the reduction in *Kcnd3* (Suppl. Figure 10).

**Figure 3:**
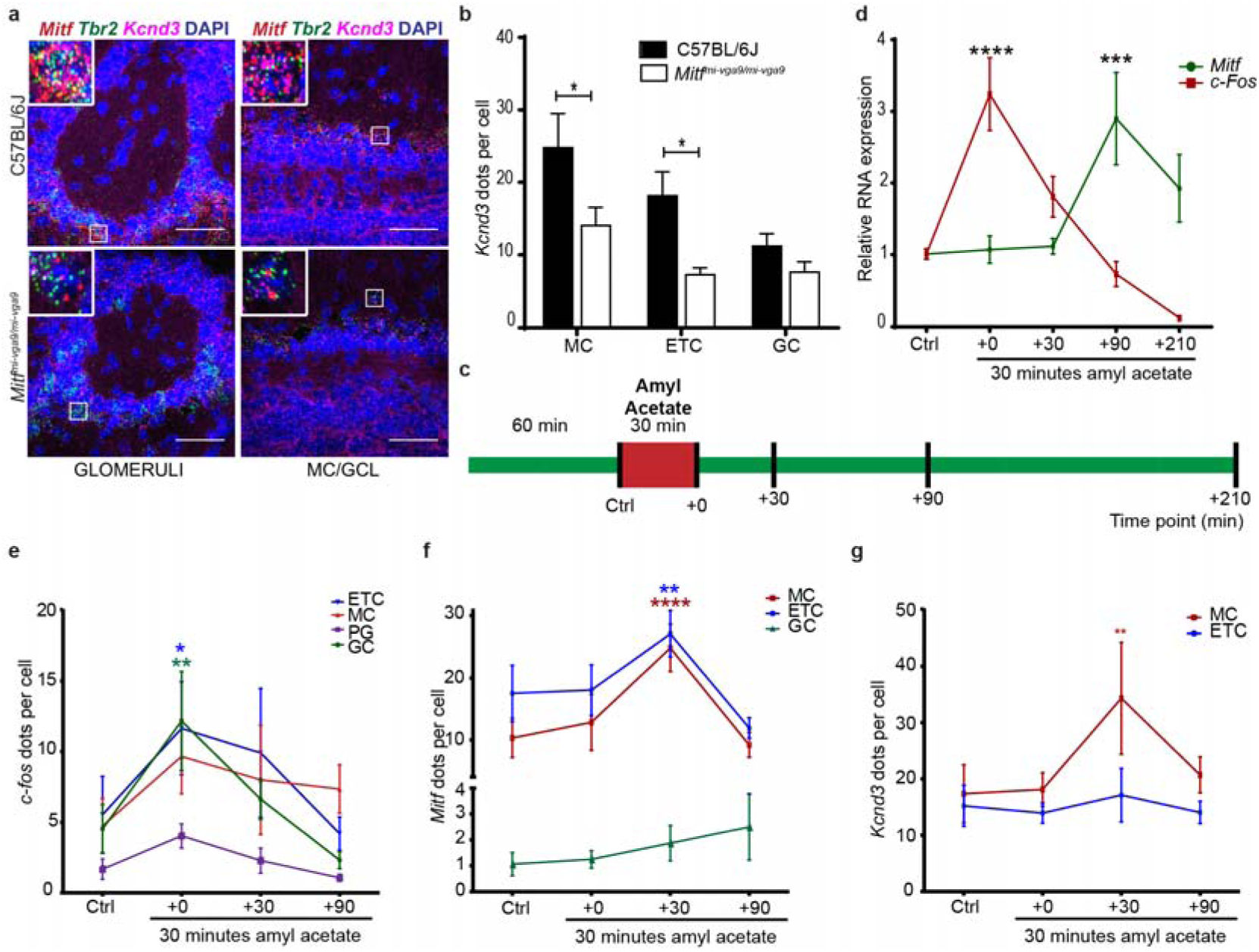
*Kcnd3* and *Mitf* expression are activity-dependent in the OB. **a.** RNA *in situ* hybridization of *Tbr2* (green), *Mitf* (red) and *Kcnd3* (magenta) performed on wild-type and *Mitf^mi-vga9/mi-vga9^* OBs. **b**. *Kcnd3* count per cell. N=5 per genotype. **c.** Schematic representation of the amyl acetate (AA) experiment where mice were habituated to an odourless cage for 60 minutes. Upon exposure to AA for 30 minutes (red block), they were sacrificed at different time points (black vertical lines) **d.** *Mitf* and *c-Fos* mRNA expression in wild-type OB following AA, determined by RT-qPCR. N=6 per time point. **e.** *c-Fos* dots per cell in wild-type OB mice following AA. N=6 per time point. **f.** *Mitf* dots per cell in wild-type OB following AA. N=6 per time point. **g**. *Kcnd3* dots per cell in wild-type OB following AA. N=6 per time point. The values on the graphs are mean ± SEM. DAPI nuclear staining is shown in blue. Scale bars are 50µm. P-values were calculated using two-way ANOVA (**b,d,f,g,i**). *P<0.05, **P<0.01, ***P<0.001, ****P<0.0001. PG= periglomerular.

Neuronal activity leads to generation of cAMP, activation of the MAPK pathway kinases and the cAMP responsive element binding protein (CREB) protein. These are established regulators of Mitf transcription and MITF signaling [37]. We therefore explored the effects of OSN activity on the expression of *Mitf* and *Kcnd3* in the OB. Amyl-acetate is known to activate a multitude of glomeruli [38]. Amyl acetate treatment of the wild type mice (Figure 3c) resulted in an immediate increase in the global transcription of *c-Fos* (Suppl. Figure 11, Figure 3e; q(40)=4.875, p<0.0001, Dunnett’s multiple comparison) as expected. Similarly, RNA *in situ* hybridization showed significant cell-specific increase of *c-Fos* in the ETCs (q(57)=2.687, p=0.0241, Dunnett’s multiple comparison) and granule cells (q(57)=3.359, p=0.0036, Dunnett’s multiple comparison) upon treatment. An increase was also observed in *c-Fos* expression in MCs, but this was not significant (q(57)=2.365, p=0.0561). Note, however that not all glomeruli are affected by amyl acetate [38]. The increase in *c-Fos* expression was followed by an increase in *Mitf* expression 30 minutes after exposure to amyl acetate both in the ETCs (Suppl. Figure 11, Figure 3f) (q(45)=3.325, p=0.005, Dunnett’s multiple comparison) and the MCs (q(45)=5.031, p=0.0001, Dunnett’s multiple comparison). Expression of *Kcnd3* also increased 30 minutes after exposure, but only in the MCs (Suppl. Figure 12, Figure 3g; q(30)=3.271, p=0.0075, Dunnet’s multiple comparison) and not in the ETCs. This showed that expression of both *Mitf* and *Kcnd3* is activity-dependent in the MC projection neurons of the OB. The large increase observed for *Mitf* expression after 90 minutes (Figure 3d), suggests that *Mitf* is also activity dependent in the granule cell layer, but the granule cells greatly outnumber M/T neurons.

### Activity-dependent increase in *Kcnd3* expression requires MITF enhancer activity

Induction of large-scale OB activity with amyl acetate was also performed on *Mitf* mutant mice. The increase observed in *Kcnd3* expression upon inducing neuronal activity in wild type OBs was not observed in *Mitf* mutant OBs (Figure 4a, b). Hence, *Mitf* and *Kcnd3* are both regulated in an activity-dependent manner in the MC and the expression of *Kcnd3* depends on *Mitf* in both MC and ETC. Analysis of two MITF ChIPseq data sets [34, 36] showed that MITF binds to intronic regions of *KCND3* in 501mel melanoma cells. The MITF ChIPseq datasets show the same overlapping binding peak B (Figure 4c, orange). A second MITF binding peak, peak A, was observed only in the Laurette dataset and overlaps with H3K27ac and H3K4me1 ChIPseq peaks in 501mel melanoma cells [39] indicative of an active enhancer (Figure 4c, yellow). Transcription activation analysis showed that MITF activates expression from a fragment containing peak B in both HEK293T cells (Figure 4e; MITF-pCMV vs pCMV: q(12)=22.58, p=0.0001; MITF-pCMV vs MITFB4RA-pCMV: q(12)=20.78, p=0.0001, Dunnett’s multiple comparison) and N2A cells (Figure 4f; MITF- pCMV vs pCMV: q(12)=15.39, p=0.0001; MITF-pCMV vs MITFB4RA-pCMV: q(12)=15.18, p=0.0001, Dunnett’s multiple comparison), whereas transcriptionally inactive MITF does not (Fig. 4 e,f). *Tyr* was used as a positive control and was significantly activated in both cell types. No activation was observed in either cell type from a fragment containing peak A. As the region around peak B has active enhancer marks, we conclude that MITF signaling is induced by activity and regulates an enhancer region in an intron of *Kcnd3* leading to an activity-dependent increase in *Kcnd3* expression in projection neurons of the OB. In accordance with this, stimulus dependent enhancer activity has previously been described in neurons [10, 40–43].

**Figure 4.**
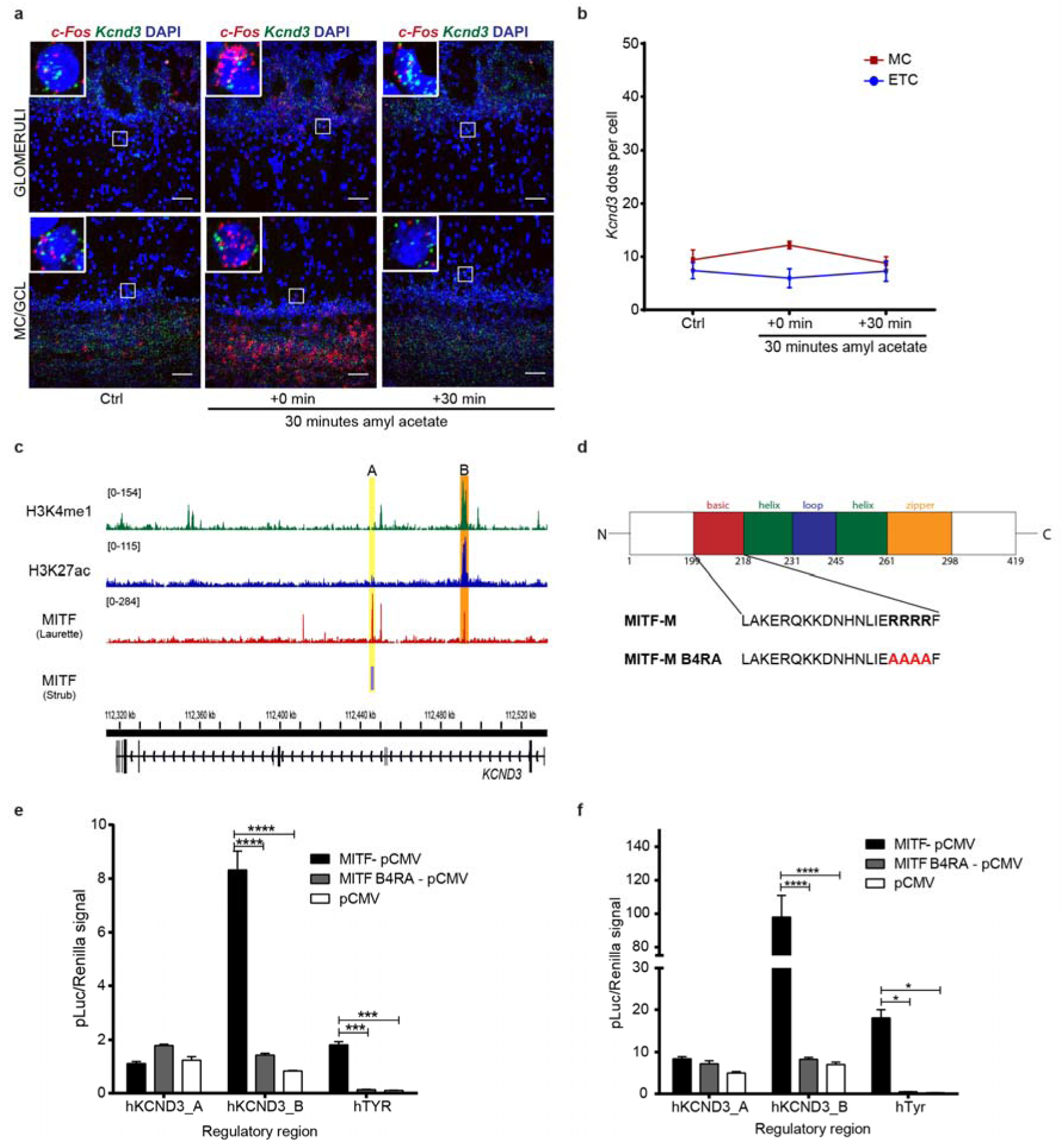
Activity-dependent increase in *Kcnd3* expression requires MITF enhancer activity. **a.** RNA *in situ* hybridization of *c-Fos* (red) and *Kcnd3* (green) performed on *Mitf^mi-vga9/mi-vga9^* OB following AA. **b**. *Kcnd3* dots per cell in *Mitf^mi-vga9/mi-vga9^* OB following AA. N=3-5 per time point. **c**. ChIPseq peaks of MITF, H3K27ac and H3K4me1 on *KCND3* gene in 501mel cells. The MITF peaks are labeled A (orange) and B (yellow). Yellow shows overlapping peak of MITF binding in Strub and Laurette datasets (B) whereas Orange indicates MITF peaks that overlap with H3K27ac and H3K4me1 peaks (A). **d.** Sequence of the basic region of wild-type (MITF-M) and transcriptionally inactive MITF with four argenines mutated to alanines (MITF-M B4RA). **e.** Transcription activation assays performed in HEK293T cells co-transfected with constructs containing the hTYR, hKCND3_pA, and hKCND3_pB regulatory regions fused to luciferase, together with empty vector or wild-type or transcriptionally inactive (B4RA) MITF constructs. N=3. **f**. Transcription activation assays performed in N2A cells co-transfected with constructs containing the hTYR, hKCND3_pA, and hKCND3_pB regulatory regions fused to luciferase, together with empty vector or wild-type or transcriptionally inactive (B4RA) MITF constructs. N=3. The values on the graphs are mean ± SEM. DAPI nuclear staining is shown in blue. Scale bars are 50µm. P-values were calculated using two-way ANOVA (**b, e, f**) *P<0.05, ****P<0.0001.

### Loss of *Mitf* affects long term adaptation

Hyperactive MCs are likely to affect olfaction. Changes in the olfactory neuronal circuits frequently lead to reduced ability to detect odorants or to differentiate between them. Both are critical functions of the olfactory system. Olfactory testing, however, showed no difference in the ability of *Mitf* mutant and wild-type mice to detect or avoid odours (Figure 5a, b, Supplementary Table 1). Next, we tested the ability of *Mitf* mutant mice to differentiate between odourants. Repeated introduction of the same odourant leads to reduced interest (habituation) and when followed by a novel odourant, renewed interest (dishabituation) (Figure 5c) - an established measurement of reduced olfactory discriminatory ability [44]. In this assay, the *Mitf* mutant mice showed increased dishabituation when exposed to the chemically similar odourant pair vanilla-almond (Figure 5d; t(84)=5.223, p<0.0001, Dunnett’s multiple comparison). This was also observed when the odourants were introduced in the reverse order (Figure 5e; t(126)=5.838, p=0.0001, Dunnett’s multiple comparison) and when the less chemically similar odourant pair lemon-vanilla was used (Figure 5f; t(147)=2.708, p=0.0144, Dunnett’s multiple comparison). This can be interpreted as increased olfactory discriminatory ability or increased olfactory sensitivity. It is also possibility that the mutant mice are more interested in the novel, second odorant. Importantly, *Mitf^mi-vga9/+^* heterozygotes, which are phenotypically similar to wild-type mice, show an intermediate phenotype (Figure 5d-f). This demonstrates that the changes in olfaction in the *Mitf* homozygotes are not due to larger cortical area devoted to olfactory processing caused by their blindness and deafness. Odour exposure reduces sensitivity of M/T neurons [15] and the appropriate long-term adaptation to sensory input is fundamental to olfaction. Our results suggest that the mutant mice might not adapt well on a longer time-scale, as *Mitf* dependent transcription would be required. We therefore tested the effects of a long-term exposure of an odourant on olfactory ability, also termed odour fatigue. Under normal circumstances this leads to a reduction in ability to detect the odorant (habituation), followed by recovery once the odorant is removed. However, in the mutant mice, there was a stark reduction in the ability to detect the odourant as 6 out of 8 mutant mice tested did not detect the initial odor when reintroduced (Figure 5g; t(84)=3, p=0.0246, Sidak’s multiple comparison). As this odorant is novel when reintroduced, this makes it unlikely that the *Mitf* mutant mice are only more interested in new odors. Both assays thus indicate clear differences in olfactory mechanisms between wild type and mutant mice.

**Figure 5:**
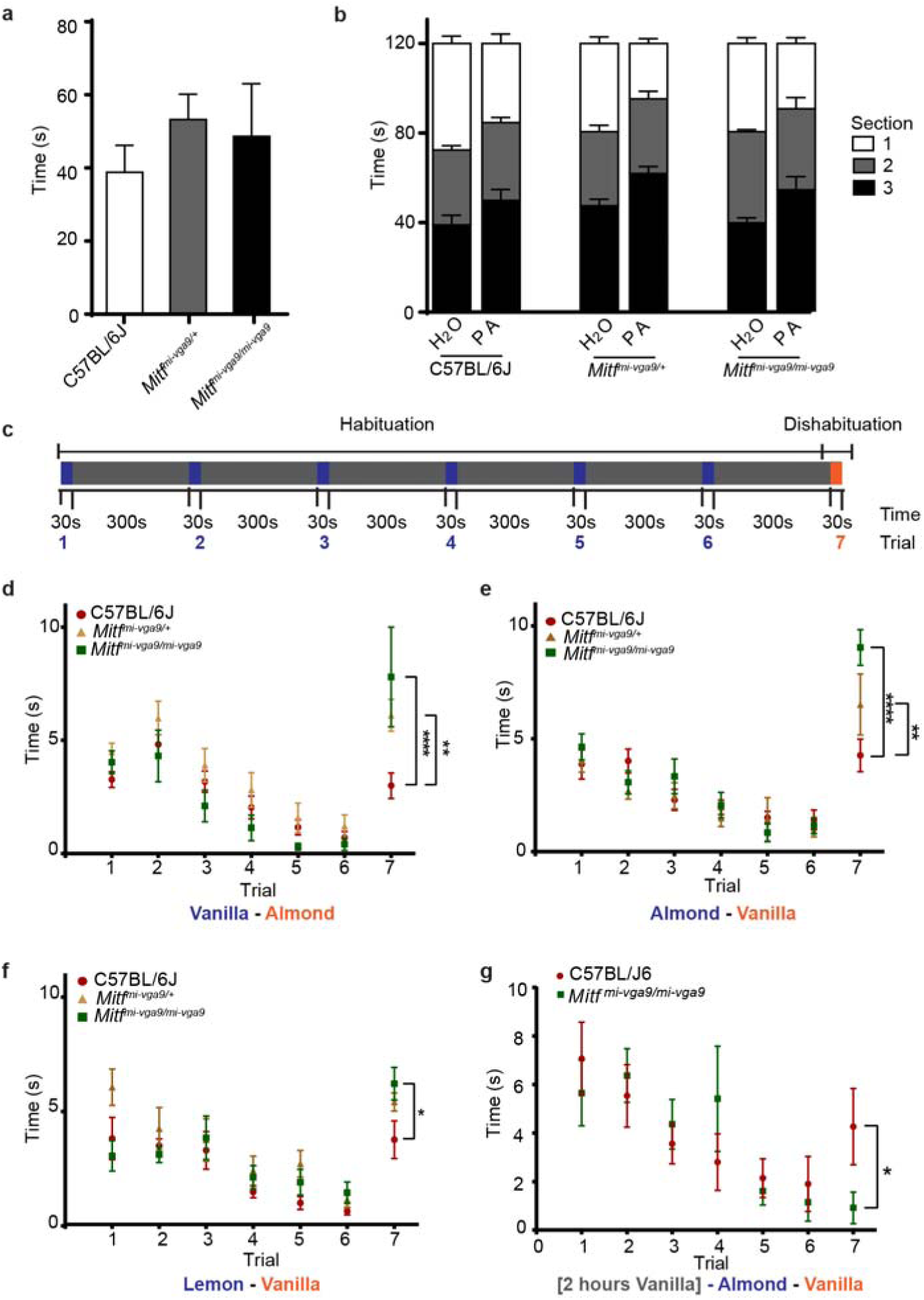
*Mitf* regulates olfactory discriminatory ability and odor fatigue recovery. **a.** Quantification of the results of the hidden cereal odour detection-assay. N=8 per genotype **b.** Avoidance assay where a cage was sectioned in 3 equal areas and mice were exposed to water or propionic acid (PA) in section 1. Quantification shows time spent in each section. Supplementary Table 2. N=6-8 per genotype. **c**. Schematic overview of the dishabituation-habituation assay, where mice are exposed to odour A (red) for 30 seconds 6 times with 5-minute intervals, after which they are exposed to odour B (green) for 30 seconds. **d** Dishabituation-habituation, mice were exposed to vanilla as odour A and almond as odour B. N=8 per genotype. **e**. Dishabituation-habituation, where mice were exposed to almond as odour A and vanilla as odour B. N=6-8 per genotype. **f**. Dishabituation-habituation, mice were exposed to almond as odour A and lime as odour B. N=8 per genotype. **g.** Dishabituation-habituation assay where the mice were exposed to vanilla for two hours, followed by almond as odour A and vanilla as odour B. N=15 per genotype. The values on the graphs are mean ± SEM. P-values were calculated using one-way ANOVA (**a**) or two-way ANOVA (**b, d-g**). *P<0.05, **P<0.01****P<0.0001.

We propose a model for a functional role of *Mitf* in projection neurons of the olfactory bulb, where it regulates the intrinsic excitability of the neurons following activity induction, by reducing the neuronal firing rate of neurons through its regulation of *Kcnd3* expression. In this model, the activity of OSNs leads to an increase in *Mitf* expression, and an *Mitf*-dependent increase in the transcription of *Kcnd3* in the output neurons of the OB. An increased I_A_- current, in turn, decreases the likelihood of the generation of action potential and reduces activity leading to decreased sensitivity of the glomeruli in question (Figure 6). Conversely, lack of activity in a glomerulus would lead to the opposite mechanism and increased sensitivity [17].

**Figure 6:**
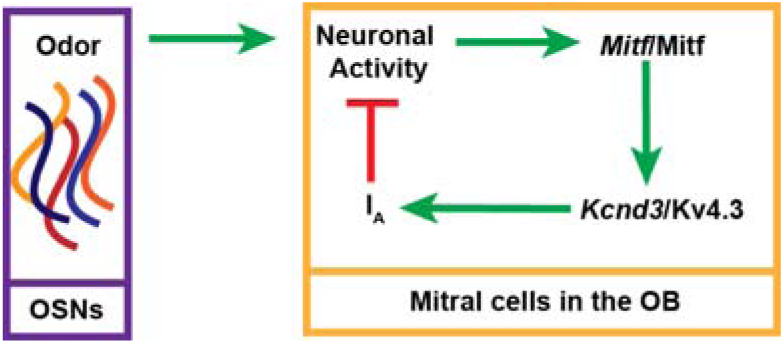
Model of MITF as a regulator of homeostatic intrinsic plasticity. Upon neuronal activity in the OB, MITF is required for an increase in *Kcnd3* expression. More Kv4.3 protein results in an increase in the I_A_ potassium current and reduced likelihood of action potentials, subsequently leading to decreased activity of M/T cells.

## Discussion

The transcription factor *Mitf* is expressed distinctly in the excitatory neurons (M/T) of the mouse OB [19]. M/T neurons are generated in the *Mitf* mutant mouse and express the appropriate neuronal markers suggesting that loss of *Mitf* does not affect their development. However, electrophysiological analysis of M/T cells *in vitro* showed that these cells are hyperactive and show loss of the potassium I_A_-current. This is further supported by reduced gene expression of *Kcnd3*, a potassium channel subunit required for the I_A_-current. Expression of the activity dependent *c-Fos* gene showed that there is an increase in neuronal activity in *Mitf* mutant mice *in vivo*. However, this was not observed in all M/T cells. External tufted cells (ETC), a subclass of tufted cells, which drive olfactory output [45] and link OSN with M/T neurons [46], do not show increased expression of *c-Fos* under baseline conditions. Mitral cells (MC), on the other hand, which are the projection neurons of the OB, showed increased expression of *c-Fos* in *Mitf* mutant OBs. Interestingly, by inducing olfactory activity in wild type mice using a strong odorant known to activate many glomeruli, we observed an increase in both *Mitf* and *Kcnd3* expression in the MCs. The expression of both *c-Fos* and *Mitf*, but not *Kcnd3*, was increased upon activity in the ETC. This demonstrates that the MCs and ETCs, which have different functions in the olfactory neuronal circuit, also have a different intrinsic response to activity.

The increased activity of projection neurons has wide-ranging consequences for OB function. The *Mitf* mutant mice show increased olfactory dishabituation, which might mean increased ability to discriminate between odours or increased olfactory sensitivity. This is a rarely reported characteristic of mutant mice [47], and the expectation is that olfaction works at maximum discriminatory power, since this is most beneficial for the organism. As we detect no changes in mobility, motivation nor interest in novelty in other assays (Figure 5a,b,g), we conclude that this phenotype is inherent to the OB, also due to the fact that the heterozygote shows an intermediate phenotype. One explanation, for this difference in dishabituation is the increased number of glomerular excitatory neurons of the OB. The nature of the olfactory phenotype in the *Mitf* mutant mouse is also likely due to increased levels of lateral inhibition due to increased activity [48, 49] and/or more glomeruli being activated by each odourant due to lower threshold of firing, leading to reduced cross-habituation [50]. Further studies will determine this. Importantly, while the *Mitf* mutant mouse has increased dihabituation, long-term odour exposure leads to a reduction in the ability to detect that odorant i.e. increased habituation. This shows that *Mitf* is required for normal long-term adaptation in the OB and that the increase in olfactory sensitivity observed in the mutant mice comes at a cost. Likely functional explanations are either that projection neurons are overwhelmed with activity or that high activity of projection neurons in the mutant leads to stronger adaptation in the olfactory cortex. Both cases show how important *Mitf* is for olfactory adaptation.

Neuronal activity sculpts the nervous system and is required for adaptive changes in neuronal function. To do so, neuronal activity must functionally change the neuron, by impacting its firing frequency or the strength of individual synapses. However, neurons also need to tightly regulate their own activity: too much neuronal activity is costly and can lead to excitotoxicity or uncontrolled firing, whereas the loss of neuronal activity leads to loss of information and neurodegeneration. To avoid this, mechanisms are in place to maintain neuronal activity within a set window, and the ability of a neuron to be plastic has to be within the confines of such neuronal homeostasis [5, 51, 52]. Our results show how sensory activity can affect firing rate of the olfactory bulb projection neurons in an *Mitf* dependent manner through effects on A-type currents via *Kcnd3* expression. This mechanism can maintain neuronal homeostasis or determine the sensitivity or neuronal gain-control of the projection neurons on a longer time scale, i.e. it is based on transcriptional changes. Importantly, electrophysiological analysis in primary neurons shows that changes in potassium currents are cell-autonomous and not due to changes in OB circuits. Detailed analysis of the electrophysiology of M/T neurons in OB slices and *in vivo* is needed to further dissect the consequences for olfaction.

Sensory systems focus on detecting novelty or change. In olfaction, this translates into the ability to quickly distinguish rare but important odours from background odours [15, 17]. In the OB, *Mitf* is pivotal to the adaptation of individual glomeruli through regulating intrinsic homeostatic plasticity in an activity-dependent manner. Most glomeruli are activated rarely by odourants. However, they can play important roles in detecting rare threats or food sources, both of high survival value. Keeping these glomeruli sensitive is therefore of high adaptive value and our model provides a mechanism through which this can occur. Thus, regulation of intrinsic plasticity, as described here, provides a mechanism through which olfaction can be insensitive to background odours but sensitive to rare odours.

## Materials & Methods

### Animals

All animal procedures were approved by the Committee on Experimental Animals, according to regulation 460/2017 and European Union Directive 2010/63 (license number 2013-03-01). Mice of the genotypes *C57BL/6J* wild-type), *C57BL/6J-Mitf^mi-vga9/mi-vga9^* and *C57BL/6J-Mitf^mi−vga9/+^* (*Mitf^mi-vga9/+^*) were maintained at the mouse facility at the University of Iceland. Unless otherwise specified, mice were housed two to five per per cage in a temperature-controlled environment (21-22°C). Unless otherwise specified, mice consumed water and food *ad libitum*. Mice of P75-P95 of both genders were used for all experiments.

### Electrophysiology

Primary OB neuronal cultures from P0-P2 mice were obtained as described by [53] with slight modifications. Briefly, mice were decapitated and the head submerged in ethanol, upon which the head was placed in Ca^2+^ and Mg^2+^-free HBSS (Gibco) containing 1% Penicillin/Streptomycin (Gibco) and 1% Amphoterecin B (Gibco) under a dissecting microscope. OBs were dissected from the heads, the meninges removed and the OBs then trypsinized for 7-10 minutes at 37°C. The cells were washed with 0.05% trypsin-EDTA prior to incubation in the trypsin solution. The neurons were then gently resuspended in Neurobasal-A Medium (Gibco) supplemented with 1% GlutaMAX (Gibco), 1% Penicillin/Streptomycin, 1% Amphoterecin B, 2% 50x serum-free B27 Supplement (Gibco) and 10% FBS. The cells were diluted to a density of 1×10^6^ cells·mL^−1^ and plated at 1.25×10^5^ cells·cm^2^ on 0.1mg/mL Poly-D-Lysine hydrobromide (Sigma-Aldrich) and 0.075mg·mL^−1^ PureCol Purfied bovine Collagen Solution, Type 1 (Advanced BioMatrix) pre-coated coverslips in Nunclon Delta Surface 24-well plates (Thermo Scientific). Neurons were cultured in Neurobasal-A Medium supplemented with 1% GlutaMAX 1% Penicillin/Streptomycin, 1% Gibco Amphoterecin B and 2% 50x serum-free B27 Supplement for 2 days, upon which they were placed in GlutaMAX-free Medium, which was replenished every 3 days.

The cultured cells were maintained in a solution with NBM-B27, 1% GlutaMAX until DIV = 2, and subsequently in NBM-B27 without GlutaMAX. The medium was replaced every 2-3 days. The mean age of cultures at the time of analysis was 12 +/− 3 days. The osmolarity of the solutions was measured with an osmometer and the pipette solution was 319 mOsm, while the extracellular Krebs solution was adjusted to 312 mOsm. Isolated cells on coverslips were placed in a recording chamber, and kept at room temperature. The cells were continuously superfused during recordings with an oxygenated extracellular solution. The extracellular mammalian Krebs solution contained the following (all in mM): 150 NaCl, 5 KCl, 2.6 CaCl_2_, 2 MgCl_2_, 10 HEPES, 5 glucose, at pH 7.3. The solution was filtered with a 0.2 µm filter prior to use. Pipettes were pulled out of 1.5mm O.D. borosilicate glass capillary tubings (Science Products GmbH) with a two-stage Narishige PP83 horizontal puller (Narishige Co), and then fire-polished to about 1 µm tip diameter in a Narishige MF-83 microforge. Pipette resistances were 2-6MΩ when placed in the Krebs solution while filled with the internal solution. The Internal solution contained the following (all in mM): 145 KCl, 2 MgCl_2_, 10 EGTA, 10 HEPES, 0.2 NaATP, pH 7.3. The reference Ag/AgCl electrode was placed in a compartment separated from the recording chamber, and electrically linked to the superfused compartment containing the cells with a glass tube bridge filled with 2M KCl/Agar solution. Recordings were made using the whole-cell configuration of the patch-clamp technique (Hamill, et al., 1981) on M/T cells with large nuclei, with an Axopatch 1D patch clamp amplifier (Axon Instruments Inc) in voltage or current clamp mode. The mean membrane resistance of wild-type M/T neurons was 104±20.3 MOhm, and 98±17 MOhm (t(25)= 0.234, p=0.82, Two-tailed test) in *Mitf* mutant neurons. The mean series resistance of wild-type M/T neurons was 13±2.3 MOhm and 17±1.6 MOhm (t(25)= −1.415, p=0.17, Two-tailed test) in *Mitf* mutant neurons. The mean cell capacitance of wild-type neurons was 46±7.3 pF and that of *Mitf* mutant neurons was 56±11 pF (t(25)= −0.7308, p=0.47, Two-tailed t-test). The mean resting membrane potentials of wild-type M/T neurons were −58±2 mV and of *Mitf* mutant neurons-57±2 mV, which is not signficantly different (t(11)=−0.102, p=0.92, Two-tailed t-test). The measured values of all these biophysical parameters of the cultured cells are comparable to previously published values measured from cultured mouse olfactory bulb M/T cells [47, 54]. The output of the amplifier was low-pass filtered at 2KHz and digitized at 10KHz with a Digidata 1440A (Molecular Devices) 16-bit A/D-D/A converter. Acquisition of recordings and generation of voltage or current pulses were performed with the pClamp 10.0 software (Molecular Devices) in conjunction with the Axopatch 1D amplifier. No corrections were made for liquid junction potentials or leak currents. After a break-in under voltage clamp, the cell was clamped at a holding potential of −70 mV, and then a 5 mV hyperpolarizing voltage step of 50 msec in duration was applied to evoke capacitance transients in the whole-cell current, to estimate the series resistance (Rs), input resistance (Ri), membrane resistance (Rm) and capacitance (Cm) of the cell. Subsequently, the cultured cells were then first voltage clamped at a holding potential of −70mV, and then stepped to −90mV for 80msec. Whole-cell outward currents were then evoked by a protocol using successive depolarizing voltage pulses from −90mV, 300msec in duration at +5 mV steps. To separate the A-type current from other outward currents another protocol with a series of voltage pulses 300 msec in duration was then applied to the cell, except that before the depolarizing pulses were delivered, the cell was held at −40mV for 80msec, after having been kept hyperpolarized for 55msec at −90mV. In order to isolate the A-type current from the total outward current, the current traces evoked by the second protocol, which primarily involves delayed rectifier K^+^ currents, were subtracted from the corresponding traces evoked by the first protocol. The resting membrane potential was recorded immediately after break in and turning to whole-cell current clamp mode with current at zero. Spontaneous membrane potential fluctuations and spiking activity of projection neurons were measured in whole-cell current clamp mode, while holding the cell near the resting potential (−65±5mV). A depolarizing shift in membrane potential larger than 25 mV was defined in the analysis software as a spike, whilst a shift of 5 mV but less than 25 mV was defined as a miniature EPSP. The time period for analysis of spontaneous spike frequency in each cell was set as 50 seconds.

### Immunofluorescence

Mice were transcardially perfused (Experimental license number 2014-07-02) with 1x Phosphate buffer solution (PBS; Dulbecco) followed by 4% paraformaldehyde (PFA; Sigma Aldrich) in 1xPBS. Following dissection, the brain was post-fixed for 2 hours in 4% PFA at room temperature. The OBs were sectioned at −20°C, in 20µm thin sections and kept at 4°C until used for immunofluorescence. Sections were placed in a blocking buffer of 0.1% Triton X-114 (Sigma Aldrich) and 5% Normal Goat Serum (NGS; Gibco) in PBS for 1 hour at room temperature. Blocking was followed by incubation with primary antibody (diluted 1:500) in blocking buffer overnight at 4°C. The antibodies used were TBR2/Eomes (Abcam, Ab23345, RRID:AB_778267) and Tyrosine hydroxylase (Millipore, MAB318, RRID:AB_2313764). Sections were subsequently incubated for 1 hour at room temperature in blocking buffer containing the Alexa Fluor goat anti-mouse 488 or Alexa Fluor goat anti-rabbit 546 (Life Technologies) secondary antibodies (each at 1:1000 dilution) and DAPI (1:1000; Sigma Aldrich). The sections were then washed and subsequently kept at 4°C until imaging at 20x magnification using an Olympus FV10-MCPSU confocal microscope; 2-5 images were obtained from the medial OB of each sample. Cells were counted manually. Quantification of the samples was performed under blind condition. The average of cells per sample was calculated and plotted using GraphPad Prism 7.

### TUNEL

Fixed frozen sections were obtained as described above. For TUNEL staining of the fixed frozen sections, In Situ Cell Death Detection Kit, Fluorescin Version 17 (Roche) was used. Images were obtained at 20x magnification using Z-stack 3D images. From each sample, 2-3 images were obtained from the medial OB. Using the Fiji software, images were stacked and positively stained cells were counted manually. The average number of positive cells per image was calculated using GraphPad Prism 7.

### Histological analysis

Mice were sacrificed by cervical dislocation and the brain was post-fixed for 24 hours in 4% Formaldehyde (Sigma Aldrich). Following fixation, the OB was embedded in paraffin and sectioned in 4µm thin sections. The sections were stained for hematoxylin and eosin (H&E).

### Plasmids

The MITF-M cDNA was cloned into the p3xFlag-CMV14 vector using EcoRI and BamHI restriction sites. The 4 arginines in the basic domain were mutated to alanines by in situ mutagenesis in order to generate the construct MITF-B4RA [55]. The human TYR promoter in pGL3 Basic Luciferase Reporter vector was used for co-transfection assays.

MITF binding sites in the intronic region of KCND3 were identified using the ChIPseq data from [34] (GSM1517751) and [36] (GSE64137). The data were imported into IGV Tools and binding peaks near KCND3 identified. Similarly, ChIPseq data for H3K4me1 (GSM2476344) and H3K27ac modifications in melanoma cells (GSM2476350) was analyzed [39]. Two MITF peaks termed A and B were identified. In order to clone the KCND3_B region, a 655bp fragment was amplified from genomic DNA of the human 501mel melanoma cells using forward primer 5‘-TTGTGAGAGTAGCAGAGTGCTTTGC-3‘ and reverse primer 5‘- GAGCAGATTCAGAGATCAGAAATCAATGG-3‘; the primers also contained restriction sites for KpnI and XhoI. In order to clone the KCND3-A region, a 469bp fragment was amplified from genomic DNA of 501mel melanoma cells using forward primer 5‘- GCTTCTGGAAGGTGAGAGAAGGA-3‘ and reverse primer 5‘- AGTGTCCTGATAGCCACATTAGGTC-3‘; the primers also contained restriction sites for NheI and XhoI (underlined). Both fragments were subsequently cloned into the pGL3 Promoter reporter vector.

### Cell culture

HEK293T and N2A neuronal cells were obtained from the Leibniz Institute DSMZ-Deutsche Sammlung von Mikroorganism un Zellkuturen GmbH. HEK293T cells were cultured in DMEM-Glutamax media (Gibco) supplemented with 10% Fetal bovine serum (FBS; Gibco), N2A cells were cultured in DMEM-Glutamax media supplemented with 10% FBS and 1% Non-Essential Amino Acids (Gibco). Cells were cultured in 75cm^2^ Nunc EasYflask with Nunclon Delta surface (Thermo Scientific) in HERAcell 240i incubator (Thermo Scientific) at 37°C and 5% CO_2_. HEK293T were passaged when 90% confluency was reached, by gently trypsinizing the cells, resuspending them in media and diluting the cells 1:10 and replating them. N2A cells were passaged when 70% confluency was reached, by gently trypsinizing the cells, resuspending them in media and diluting the cells 1:5 and replating them.

### Co-transfection assay

N2A and HEK293T cells were plated in Falcon 96-well cell culture plates at 13.800 and 20.000 cells per well respectively. After 24 hours, the N2A and HEK293T cells were transfected with 0.1 µg of DNA mixed with Fugene at 7:1 and 2.8:1 Fugene HD to DNA ratios, respectively. This mixture contained equal amounts of the luciferase reporter plasmid, renilla reporter plasmid and MITF-FlagpCMV, MITF-B4RAFlagpCMV, or FlagpCMV. Each combination was plated and transfected in triplicate. Cells were harvested after 24 hours of incubation using Dual-Glo Luciferase Assay System (Promega Corporation) as described and the luciferase and renilla luminescence measured separately on MODULUS II TURNER (Biosystems). To quantify the normalized luminescence, the luciferase and renilla luminescence values were divided by one another for each well. Subsequently an average was obtained for the triplicate. These values were then plotted using GraphPad Prism 7.

### Odour exposure

Mice were placed in empty cages, devoid of bedding, food and water for one hour in an odour free room. After one hour, mice were moved to a different room where an open eppendorf tube containing 60µL of amyl acetate (Sigma-Aldrich) was taped inside their cage at nose-level. After 30 minutes, the mice were placed in a new cage without amyl acetate and moved back to the odour free room. The mice were sacrificed by cervical dislocation at different intervals: before amyl acetate exposure, immediately after exposure and 30, 90, and 210 minutes after exposure to amyl acetate. The OBs were either flash frozen in liquid nitrogen for quantitative real time PCR or placed in OCT medium (Sakura) in a 15×15×15mm Tissue-Tek Cryomold (Sakura) and subsequently flash frozen for sectioning and RNA *in situ* hybridization.

### RNA mFISH

Mice were sacrificed by cervical dislocation. The OB was dissected out and placed in OCT compound in a 15×15×15mm Tissue-Tek Cryomold and flash frozen with liquid nitrogen. The caudal OB was sequentially sectioned unto slides, with 2 sections per slide, at a thickness of 20µm. The slides were stored in a −80°C freezer in an airtight bag until use. The samples were pre-treated using the protocol “RNAscope Sample Preparation and Pretreatment Guide for Fresh Frozen Tissue (Manual RNAScope assay)” (Advanced Cell Diagnostics). mFISH was subsequently performed according to the manufacturer’s instruction for the RNAscope Fluorescent Multiplex Kit (Advanced Cell Diagnostics) with probes 422501 (*Mitf*), 429641- C2 (*Tbr2/Eomes*), 452598-C3 (*Kcnd3*) and 316921-C2 (*c-Fos*). Sections were kept at 4°C until imaging using Z-stack 3D images with the Olympus FluoView FV10 confocal microscope at 30x magnification. Two images were obtained from the medial OB per sample - one image of the glomerular region of the OB and a separate image of the mitral and granule cell layer. Using the Fiji software, fluorescent dots per cell were counted, by encircling the cloud of *Tbr2* dots and using this area to count the fluorescent dots of interest. *Tbr2*+ cells located directly below the glomeruli, were considered external tufted cells. Similarly, *Tbr2*+ cells located in the mitral cell layer, and not in the external plexiform layer, were considered as mitral cells. In the cases where *Tbr2* coexpression was not determined, *Mitf* expression, localization and nuclear morphology was used to determine cell type. Average dots per cell was calculated for each sample and plotted using GraphPad Prism 7.

### Quantitative PCR

Mice were sacrificed by cervical dislocation and OBs flash frozen in liquid nitrogen. The samples were kept at −80°C until RNA was isolated using NucleoSpin RNA (Machery Nagel).

In order to generate cDNA, the SuperScript II Reverse Transcriptase kit (Invitrogen) was used, using 1 ug of RNA and oligo(T) primers. Mouse specific exon-spanning primers were designed using Primer-blast. The primers-pairs used were the following: *Gapdh* forward 5‘- ATGACATCAAGAAGGTGGTG-3‘, reverse 5‘-CATACCAGGAAATGAGCTTG-3‘; *Actin* forward 5‘-CACTGTCGAGTCGCGTCC-3‘, reverse 5‘-TCATCCATGGCGAACTGGTG-3‘; *Mitf* forward 5‘-AGCAAGAGCATTGGCTAAAGA-3‘, reverse 5‘- GCATGTCTGGATCATTTGACT-3‘and *c-Fos* forward 5‘- TTTCAACGCCGACTACGAGG-3‘, reverse 5‘-TCTGCGCAAAAGTCCTGTGT-3‘. Quantitative qPCR was performed using Power SYBR Green PCR Master Mix (ThermoFisher Scientific). The 2^−ΔΔ^*^C^*^t^ method was used to normalize expression to *C57BL/6J*, relative to *Gapdh* and *Actin* as described previously [56].

### Behavior

Prior to performing any behavior experiments, individual mice were placed in separate cages for at least 24 hours. Mice were used for a maximum of two behavioral experiments but only once for the same behavioral test. Each mouse was only used once for each habituation/dishabituation experiment. All behavior experiments were performed at 9pm, 1 hour after the lights were turned off. All behavior was filmed, and the behavior of interest measured as a function of time and plotted using GraphPad Prism 7.

### Hidden food assay

The food pellets were replaced by sweetened cereal – Cocoa Puffs (Nesquik General Mills) for 12 hours. After 12 hours, all Cocoa Puffs pellets were removed from the food section of the cage and one Cocoa Puffs pellet was placed into the cage. Following overnight starvation (license number 2016-05-01), a cage was filled with 4cm of bedding and a Cocoa Puffs pellet was hidden in one corner, under the bedding. The mouse was placed in the posterior section of the cage, opposite to the pellet, and the time it took the mouse to find the pellet was determined. Mice which showed obvious distress (tremors, unsteady gait or unwillingness to explore) and could not find the pellet in under 180 seconds were removed from the experiment.

### Avoidance assay

A 41.5 × 24 × 18.5cm cage was sectioned into three equal sections numbered 1-3 and the bottom covered with 0.5cm of bedding. Prior to the experiment, 30µL of water was placed onto a 5 × 5 cm piece of Whatman paper which was then taped onto section 1, at nose height. The mouse was gently placed in section 3 and allowed to roam for 120 seconds in the cage. After 120 seconds, the mouse was removed from the cage and the 5 × 5cm Whatman paper containing 30µL of water was replaced with a 5 × 5cm paper of Whatman paper containing 30µL of propionic acid (Sigma Aldrich). The mouse was gently placed in section 3 and allowed to roam for 120 seconds in the cage. After 120 seconds, the mouse was removed from the cage. The time the mouse spent in each of the 3 sections, using the nose as the reference point, was measured for both conditions.

### Habituation-Dishabituation

In order to perform the habituation-dishabituation experiment, we used the protocol of Lehmkuhl and associates [57]. Mice were kept in individual cages for 24 hours. Food and water were subsequently removed, and the mice were kept in an odour free room for 1 hour prior to the experiment. The mice were habituated for 30 seconds every 5 minutes by placing 5µL of a particular odour in the cage, a total of 6 times, and subsequently exposed to 5µL of another odour for 30 seconds. Time spent sniffing the object, or in the direction of the object was determined. Odourants used were almond, lime (brand name Dr Oetker) and vanilla extracts (brand name Katla). For long-term exposure to odour, the mice were exposed to vanilla odourant for 2 hours prior to habituation-dishabituation with almond-vanilla odour pairs.

### Statistical analysis

No statistical methods were used to predetermine sample size. Grouped analyses were used for the *in vivo* experiments as at least three mice per genotype were used. Quantitative results were analysed by one-or two-way analyses of variance (ANOVA), and two-sided unpaired or paired Student’s t-tests and Chi-square tests using GraphPad Prism 7. To obtain p-values, multiple comaparisons for ANOVA tests were performed with Livak’s correction or Dunnett’s correction in cases where there was a comparison to a control time point. All numerical results are presented as mean and SEM unless stated otherwise. Degrees of freedom are indicated between brackets.

## Acknowledgements

This research was supported by grants of the Icelandic Research Fund, Rannís grant numbers: [152715-053 and 163068-051]. We thank our colleagues from the BioMedical Center at the University of Iceland, Erna Magnúsdóttir PhD, Ragnhildur Káradóttir PhD, Margrét Helga Ögmundsdóttir PhD, Ramile Dilshat, Kimberley Anderson, Alba Sabate and Unnur Diljá Teitsdóttir for their help.

## Author Contributions

D.A.M.A. took part in designing experiments with P.H.P. and E.S., performed all experiments and analyzed the data, aside from the electrophysiology experiments. A.P provided data. T.E and H.R. performed the electrophysiology experiments. P.H.P., E.S. and D.A.M.A wrote the manuscript and T.E. commented on the manuscript at all stages.

---

**Figure 7:**
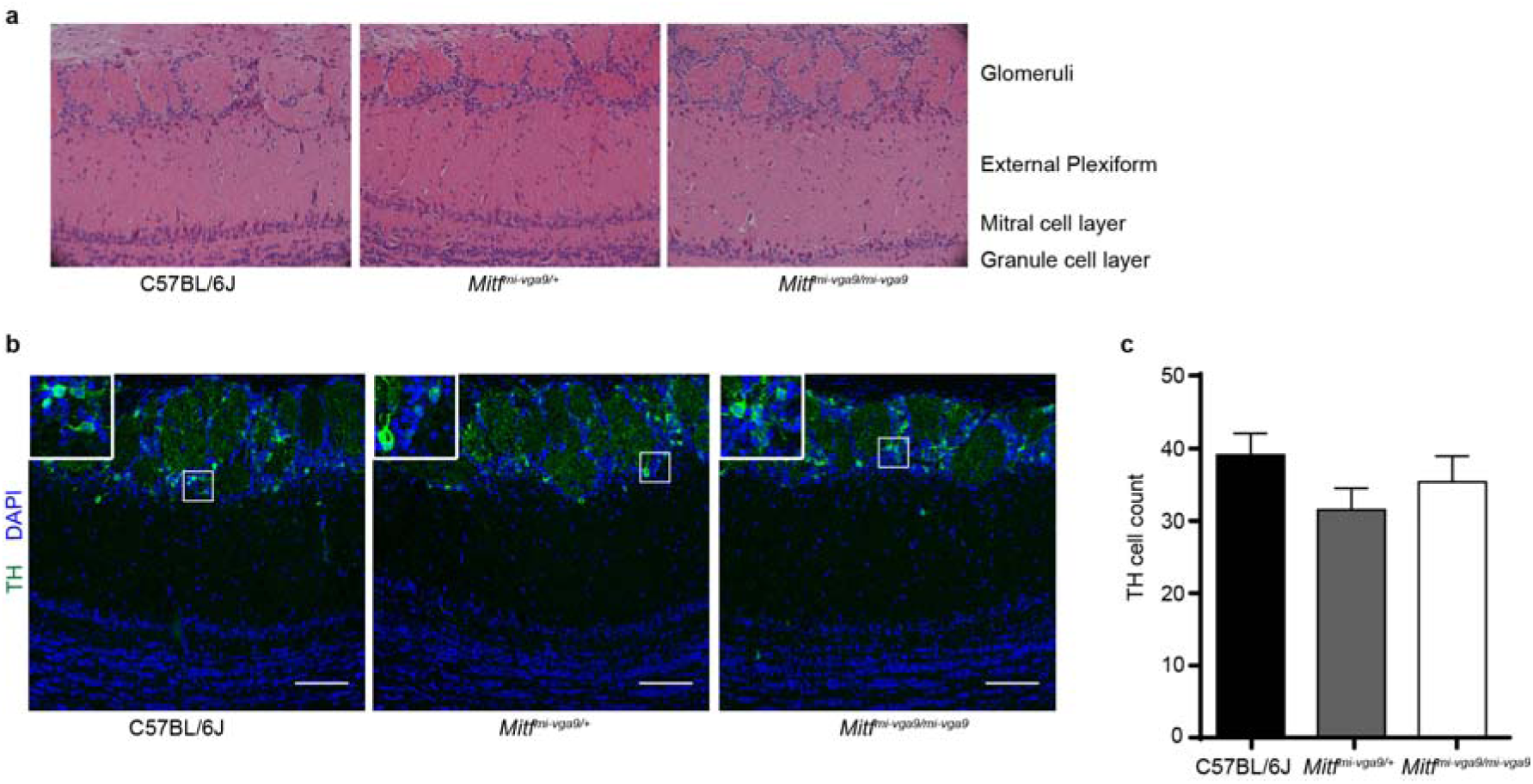
*Mitf^mi-vga9/mi-vga9^* mice have normal olfactory bulbs. **a.** Representative images of the H&E histological analysis of coronal sections of the medial olfactory bulbs of the indicated genotypes. **b.** Immunofluorescent staining of Tyrosine hydroxylase (TH) showing the localization of TH in the glomeruli of the indicated genotypes. **c.** Cell count of TH+ cells of the OB of wild-type, *Mitf^mi-vga9/+^* and *Mitf^mi-vga9/mi-vga9^* mice. N=6 per genotype. The values on the graphs are mean ± SEM. DAPI nuclear staining is shown in blue. Scale bar indicates 100µm. P-values were calculated using one-way ANOVA (**c**).

**Figure 8:**
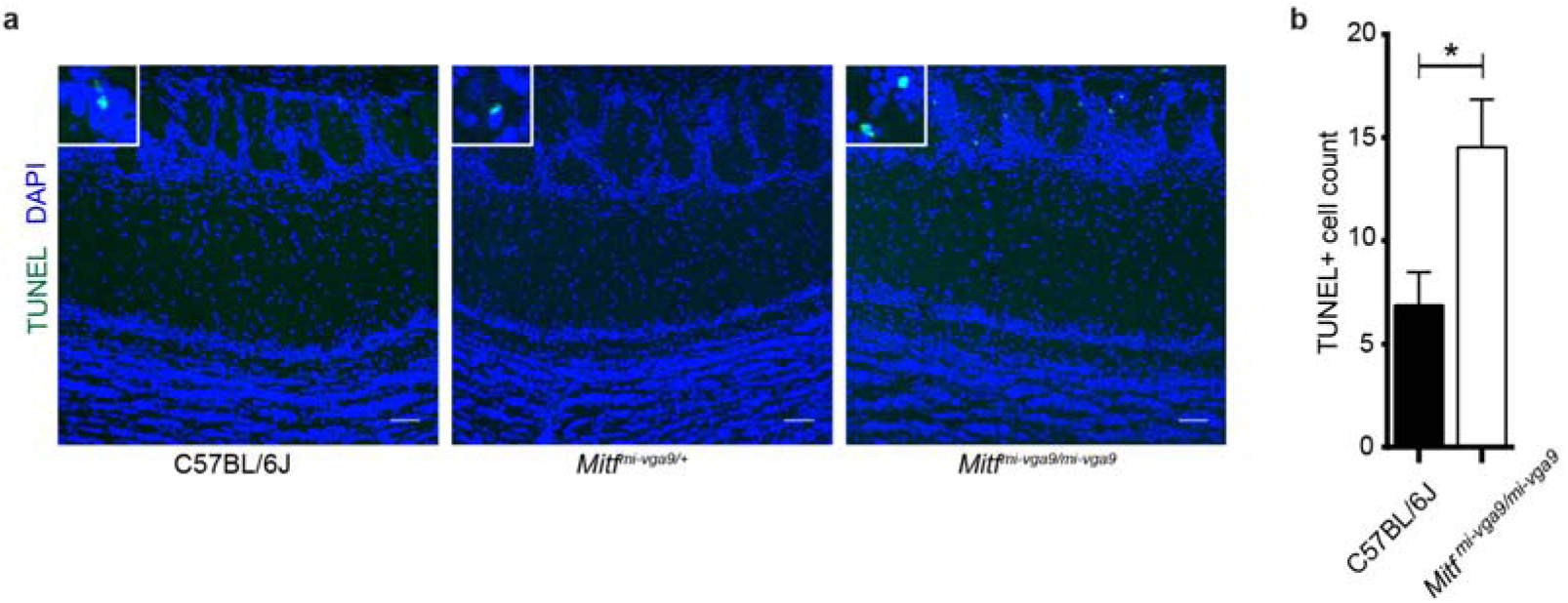
Increase in apoptosis in OBs of *Mitf^mi-vga9/mi-vga9^* mice. **a.** TUNEL staining of sections of OBs of the indicated genotypes, showing localization of apoptotic cells. **b.** The number of apoptotic cells of OB of wild-type and *Mitf^mi-vga9/mi-vga9^*mice. N=9 per genotype. The values on the graphs are mean ± SEM. DAPI nuclear staining is shown in blue. Scale bars are 50µm. P-values were calculated using two-tailed unpaired student’s t-test. *P<0.05.

**Figure 9:**
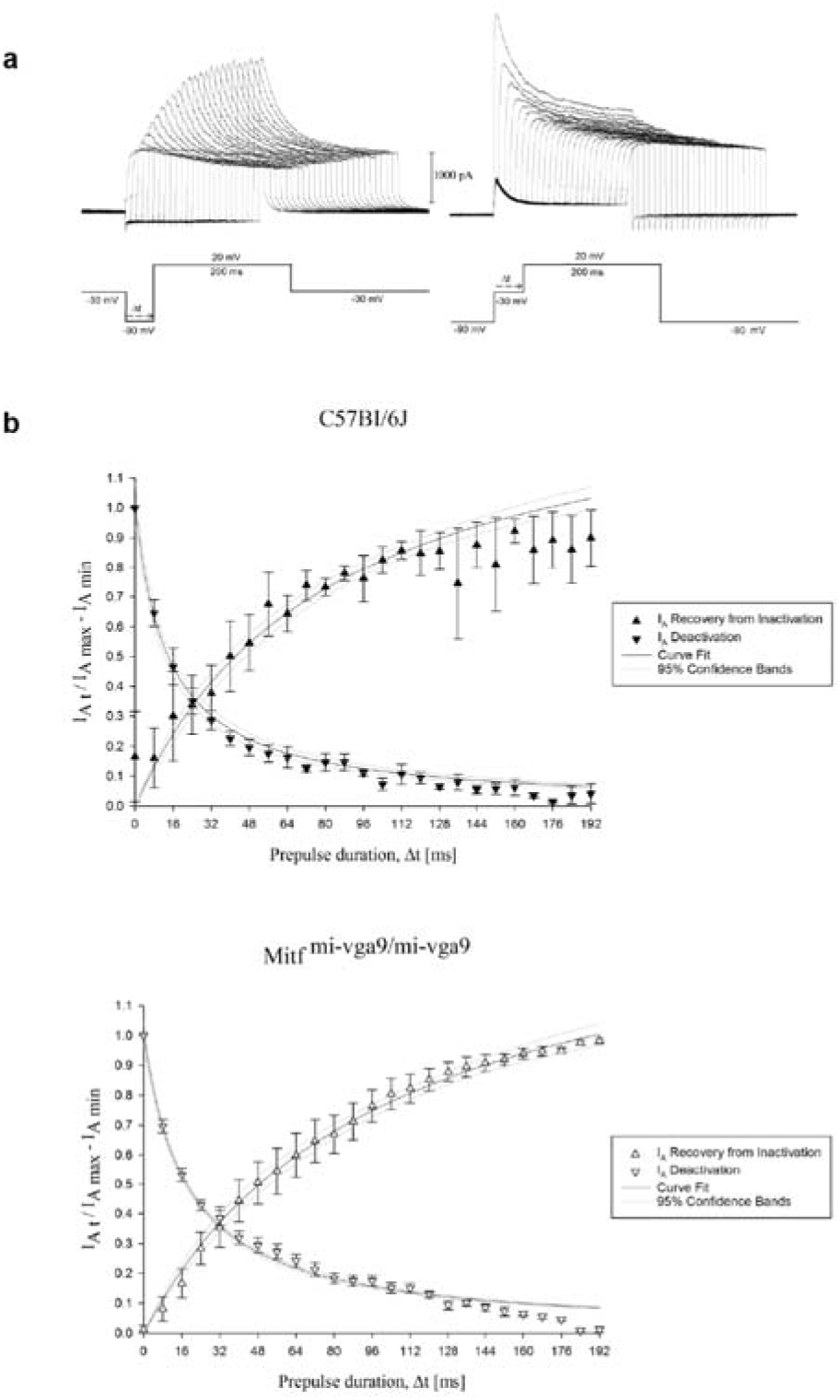
Gating kinetics of the A-type current (I_A_). **a.** Representative recordings obtained to assess the gating kinetics of the I_A_ current mediating channels. The projection neurons were first voltage clamped at −30mV and subsequently depolarized to 20mV utilizing a pre-hyperpolarizing pulse in between to recover the I_A_ current from inactivation. This was done by increasing the time, Δt, spent in the hyperpolarized state, as indicated by the voltage trace on the left. To gauge its activation kinetics, the neurons were held at −90mV, as shown in the voltage trace on the right, and subsequently depolarized to 20mV, with a pre-depolarization pulse to −30mV applied in between. The A-type current gets inactivated by increasing the duration of the pre-pulse, Δt. **b**. Normalized mean currents (±SEM) plotted separately during recovery of the I_A_ current from inactivation and during deactivation, in cells from WILD-TYPE and *Mitf^mi-vga9/mi-vga9^*mice. The functions fitted to the data yield half-activation and half-recovery times of 15.2±4.7msec and 42.6±7.2msec for the wild-type (N=5, P=0.53) and 18.2±1.5msec and 51.3±9.3msec for the Mitf^mi-vga9/mi-vga9^ (N=6, P=0.49) M/T

**Figure 10:**
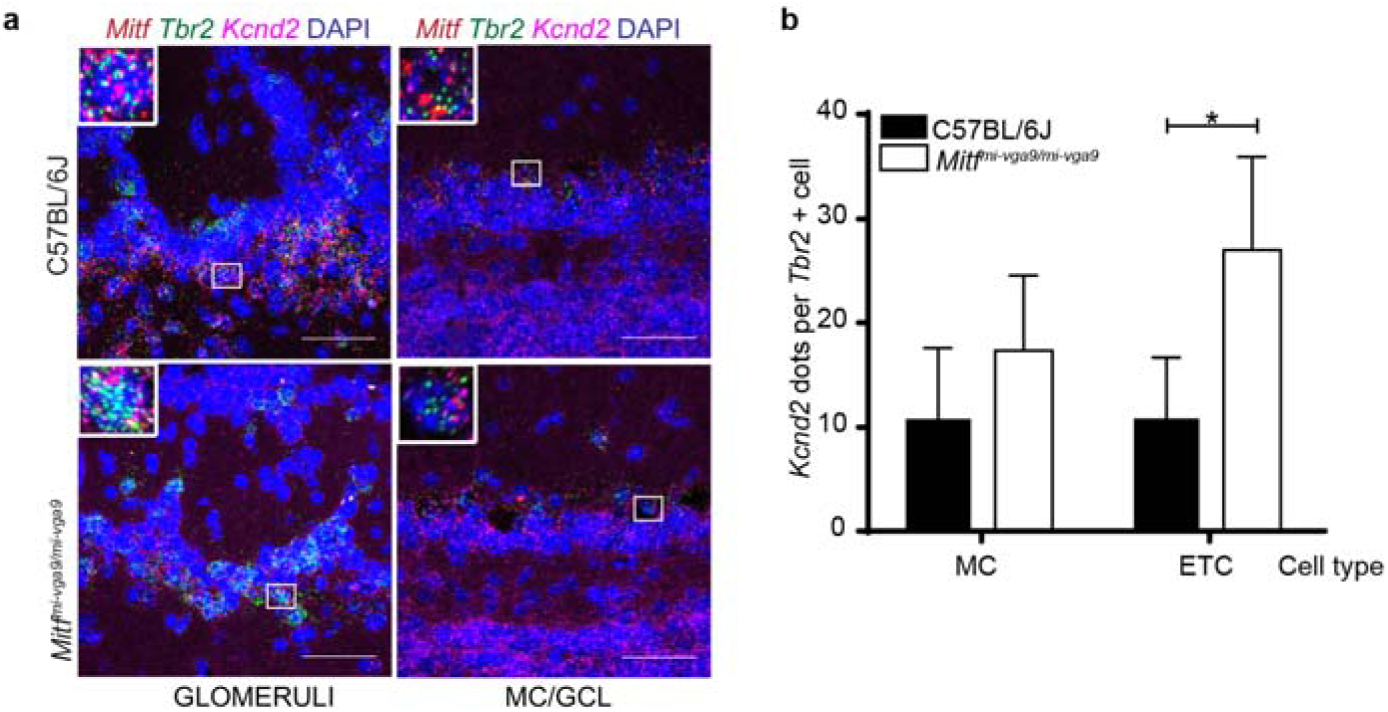
*Kcnd2* expression is increased in the *Mitf* mutant olfactory bulb. **a.** RNA *in situ* hybridization of *Tbr2* (green), *Mitf* (red) and *Kcnd2* (magenta) performed on wild-type and *Mitf^mi-vga9/mi-vga9^* OBs. **b**. *Kcnd3* count per cell. N=4 per genotype. The values on the graphs are mean ± SEM. DAPI nuclear staining is shown in blue. Scale bars are 50µm. P-values were calculated using two-way ANOVA (**b**) *P<0.05.

**Figure 11:**
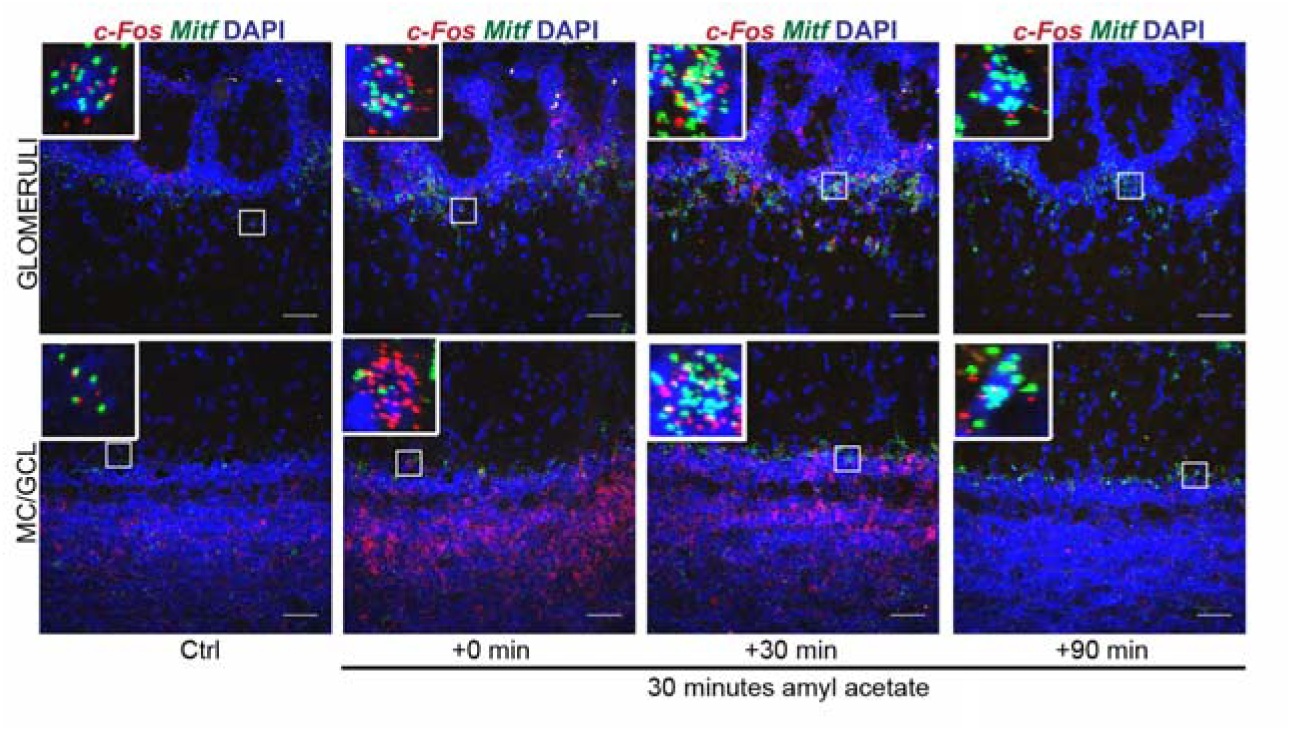
*c-Fos* and *Mitf* expression is increased upon activity induction in the olfactory bulb. RNA *in situ* hybridization of *c-Fos* (red) and *Mitf* (green) performed on wild-type OBs following AA. DAPI nuclear staining is shown in blue. Scale bars are 50µm.

**Figure 12:**
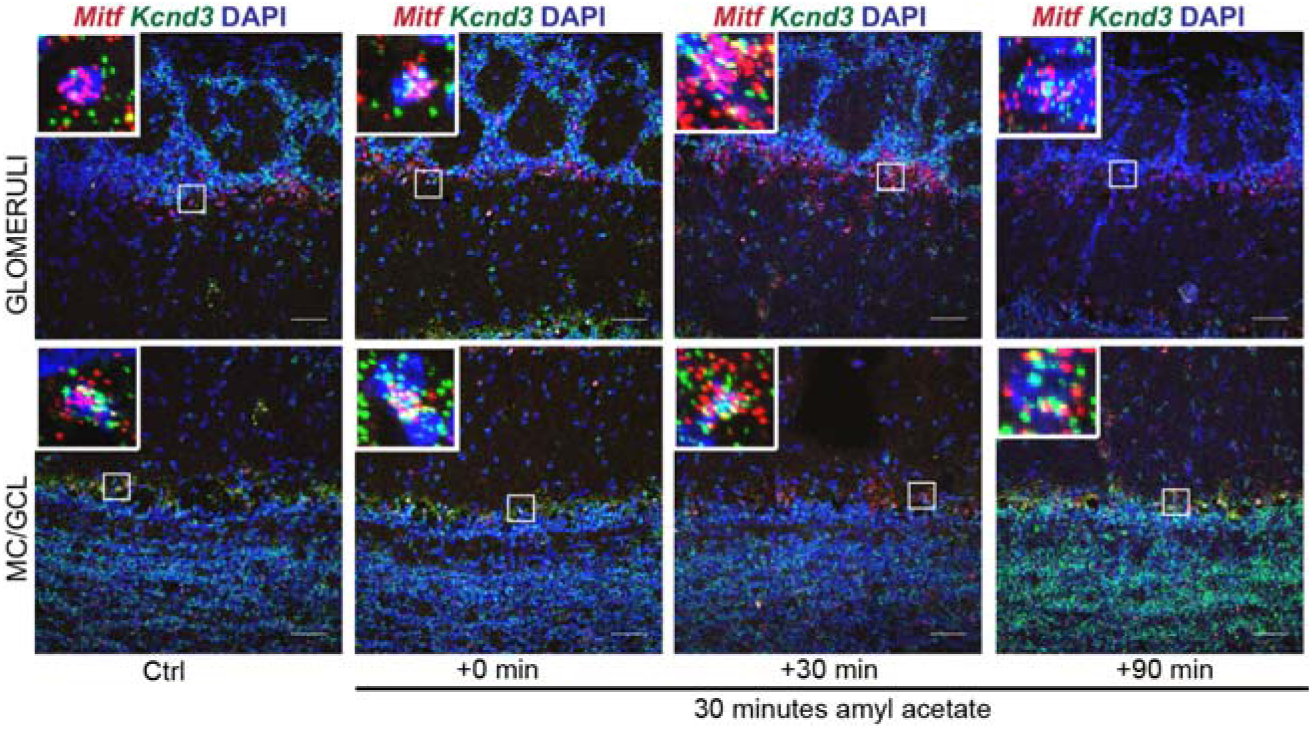
Kcnd3 expression is increased upon activity induction in the olfactory bulb. RNA *in situ* hybridization of *Mitf* (red) and *Kcnd3* (green) performed on wild-type OBs following AA. DAPI nuclear staining is shown in blue. Scale bars are 50µm.

**Table 1:**
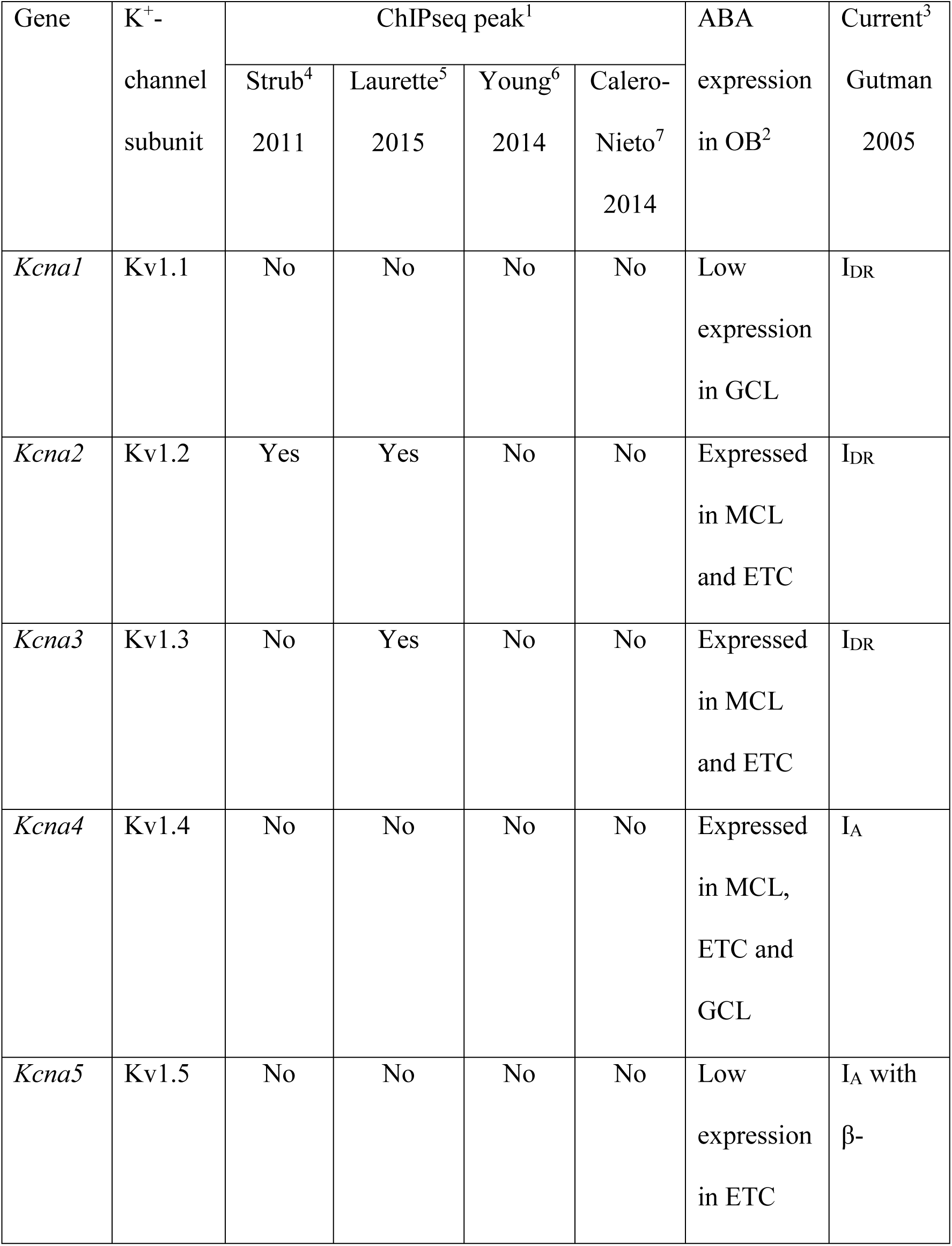
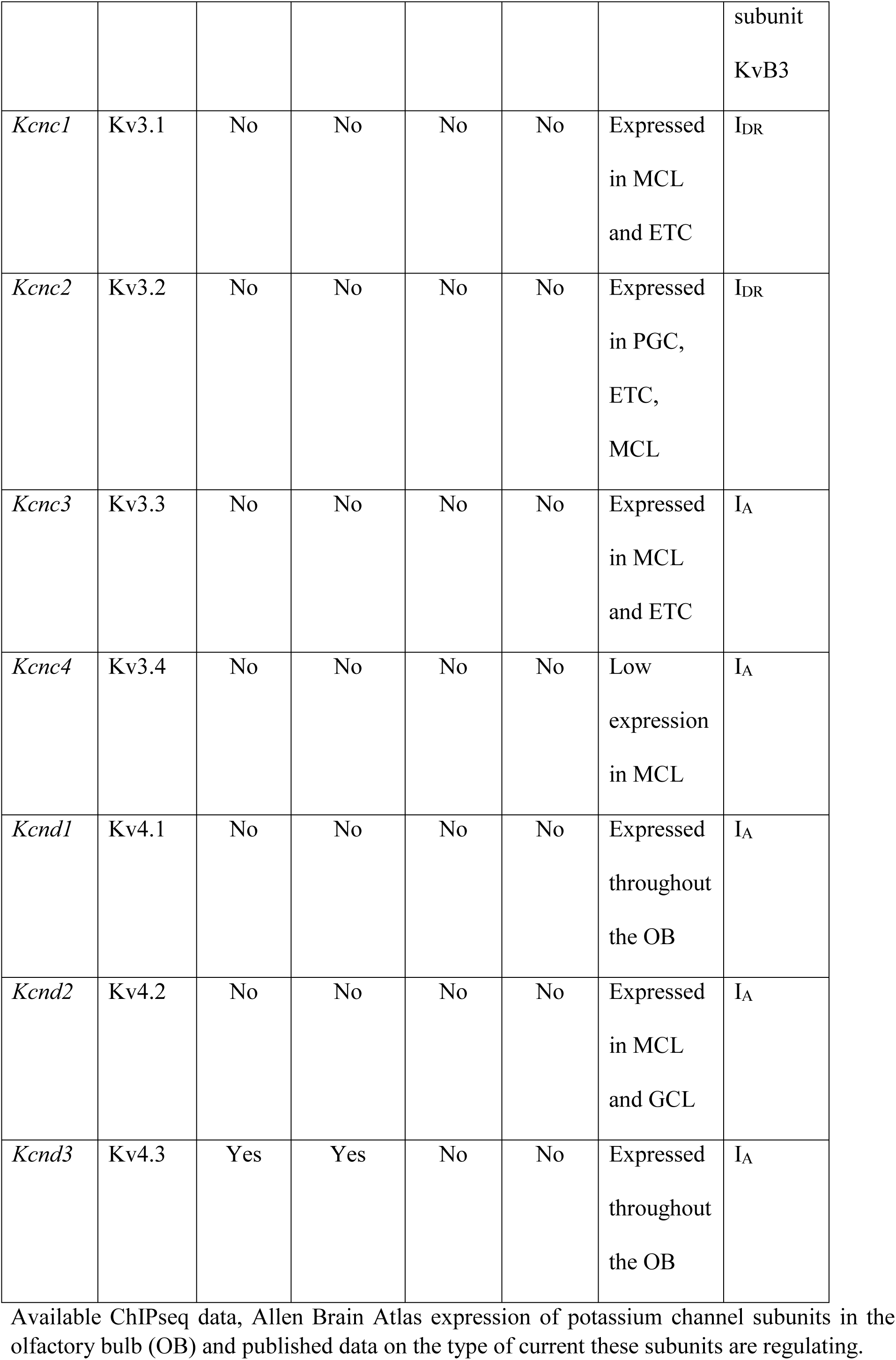
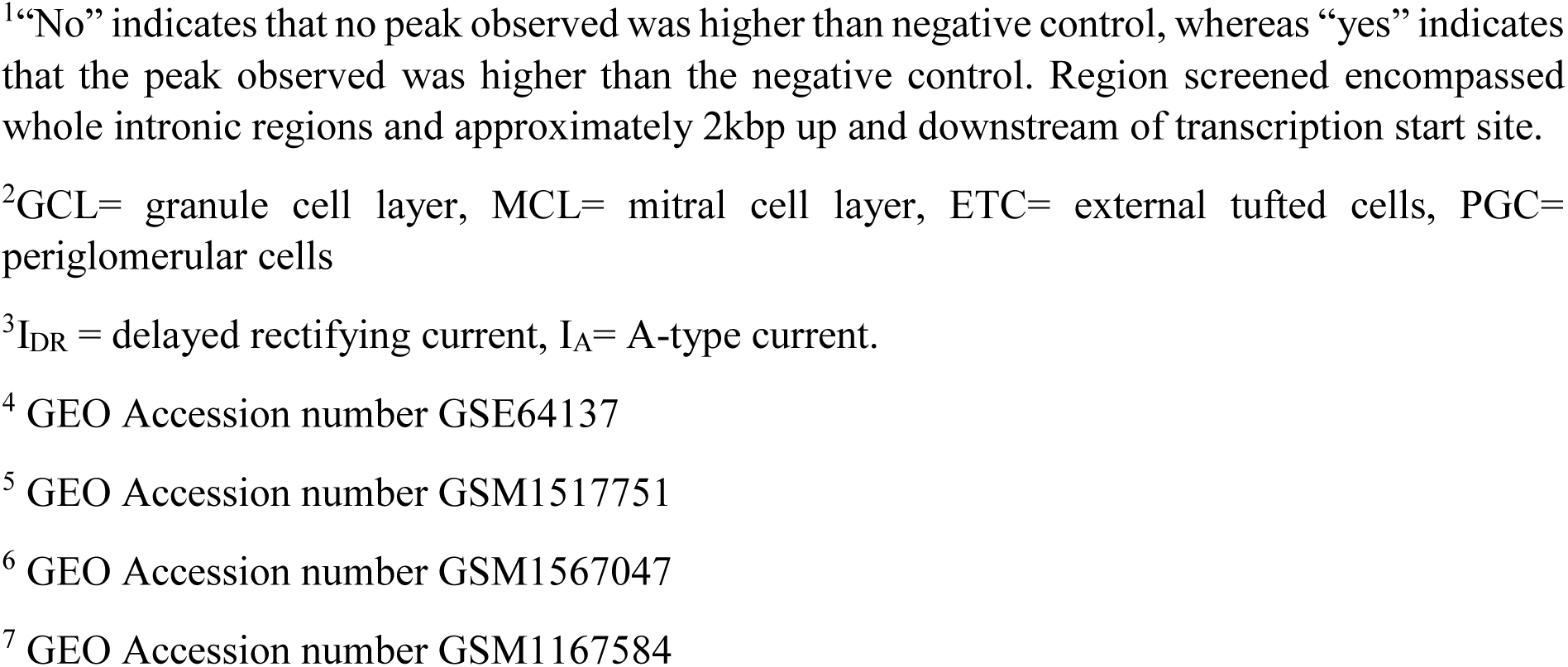
Potassium channel subunits in the OB.

**Table 2:** Two-tailed paired student’s t-test performed on avoidance behavior.

|  | C57Bl/6J | *Mitf*<sup>mi-vga9/+</sup> | *Mitf*<sup>mi-vga9/mi-vga9</sup> |
| --- | --- | --- | --- |
| Section 1 | 0.0016* | 0.0009* | 0.0004* |
| Section 2 | 0.6943 | 0.9325 | 0.3859 |
| Section 3 | 0.0408* | 0.0138* | 0.0267* |

Two-tailed paired student's t-test performed on the time mice spent in each section, comparing water and propionic acid conditions. \*P<0.05. N=6-8.

## References

1. Sweatt, J.D., Neural plasticity and behavior - sixty years of conceptual advances. J Neurochem, 2016. 139 Suppl 2: p. 179–199.

2. Yap, E.L. and M.E. Greenberg, Activity-Regulated Transcription: Bridging the Gap between Neural Activity and Behavior. Neuron, 2018. 100(2): p. 330–348.

3. Davis, G.W., Homeostatic control of neural activity: From phenomenology to molecular design. Annual Review of Neuroscience, 2006. 29: p. 307–323.

4. Turrigiano, G., Too many cooks? Intrinsic and synaptic homeostatic mechanisms in cortical circuit refinement. Annu Rev Neurosci, 2011. 34: p. 89–103.

5. Turrigiano, G.G., The self-tuning neuron: synaptic scaling of excitatory synapses. Cell, 2008. 135(3): p. 422–35.

6. Lemasson, G., E. Marder, and L.F. Abbott, Activity-Dependent Regulation of Conductances in Model Neurons. Science, 1993. 259(5103): p. 1915–1917.

7. Lisman, J., et al., Memory formation depends on both synapse-specific modifications of synaptic strength and cell-specific increases in excitability. Nature Neuroscience, 2018. 21(3): p. 309–314.

8. Kourrich, S., D.J. Calu, and A. Bonci, Intrinsic plasticity: an emerging player in addiction. Nat Rev Neurosci, 2015. 16(3): p. 173–84.

9. Hrvatin, S., et al., Single-cell analysis of experience-dependent transcriptomic states in the mouse visual cortex. Nature Neuroscience, 2018. 21(1): p. 120–+.

10. Guzman-Karlsson, M.C., et al., Transcriptional and epigenetic regulation of Hebbian and non-Hebbian plasticity. Neuropharmacology, 2014. 80: p. 3–17.

11. Kikuta, S., et al., Odorant response properties of individual neurons in an olfactory glomerular module. Neuron, 2013. 77(6): p. 1122–35.

12. Tan, J., et al., Odor information processing by the olfactory bulb analyzed in gene-targeted mice. Neuron, 2010. 65(6): p. 912–26.

13. Roland, B., et al., Massive normalization of olfactory bulb output in mice with a ‘monoclonal nose’. Elife, 2016. 5.

14. Chaudhury, D., et al., Olfactory bulb habituation to odor stimuli. Behav Neurosci, 2010. 124(4): p. 490–9.

15. Kato, H.K., et al., Dynamic sensory representations in the olfactory bulb: modulation by wakefulness and experience. Neuron, 2012. 76(5): p. 962–75.

16. Angelo, K., et al., A biophysical signature of network affiliation and sensory processing in mitral cells. Nature, 2012. 488(7411): p. 375–8.

17. Tyler, W.J., et al., Experience-dependent modification of primary sensory synapses in the mammalian olfactory bulb. J Neurosci, 2007. 27(35): p. 9427–38.

18. Tachibana, M., et al., Ectopic expression of MITF, a gene for Waardenburg syndrome type 2, converts fibroblasts to cells with melanocyte characteristics. Nat Genet, 1996. 14(1): p. 50–4.

19. Ohba, K., et al., Microphthalmia-associated transcription factor is expressed in projection neurons of the mouse olfactory bulb. Genes to Cells, 2015. 20(12): p. 1088–1102.

20. De Saint Jan, D., et al., External tufted cells drive the output of olfactory bulb glomeruli. J Neurosci, 2009. 29(7): p. 2043–52.

21. Hayar, A., et al., Olfactory bulb glomeruli: External tufted cells intrinsically burst at theta frequency and are entrained by patterned olfactory input. Journal of Neuroscience, 2004. 24(5): p. 1190–1199.

22. Whitesell, J.D., et al., Interglomerular lateral inhibition targeted on external tufted cells in the olfactory bulb. J Neurosci, 2013. 33(4): p. 1552–63.

23. Wang, F., et al., RNAscope: a novel in situ RNA analysis platform for formalin-fixed, paraffin-embedded tissues. J Mol Diagn, 2012. 14(1): p. 22–9.

24. Hodgkinson, C.A., et al., Mutations at the Mouse Microphthalmia Locus Are Associated with Defects in a Gene Encoding a Novel Basic-Helix-Loop-Helix-Zipper Protein. Cell, 1993. 74(2): p. 395–404.

25. Liu, N., et al., Unique regulation of immediate early gene and tyrosine hydroxylase expression in the odor-deprived mouse olfactory bulb. J Biol Chem, 1999. 274(5): p. 3042–7.

26. Imamura, F. and C.A. Greer, Pax6 regulates Tbr1 and Tbr2 expressions in olfactory bulb mitral cells. Molecular and Cellular Neuroscience, 2013. 54: p. 58–70.

27. Mizuguchi, R., et al., Tbr2 Deficiency in Mitral and Tufted Cells Disrupts Excitatory-Inhibitory Balance of Neural Circuitry in the Mouse Olfactory Bulb (June 27, 8831, 2012). Journal of Neuroscience, 2012. 32(39): p. 13639–13639.

28. Johnson, M.C., et al., Odor enrichment sculpts the abundance of olfactory bulb mitral cells. Neurosci Lett, 2013. 541: p. 173–8.

29. Liu, A., S. Savya, and N.N. Urban, Early Odorant Exposure Increases the Number of Mitral and Tufted Cells Associated with a Single Glomerulus. J Neurosci, 2016. 36(46): p. 11646–11653.

30. Fransen, E. and J. Tigerholm, Role of A-type potassium currents in excitability, network synchronicity, and epilepsy. Hippocampus, 2010. 20(7): p. 877–87.

31. Yuan, W., A. Burkhalter, and J.M. Nerbonne, Functional role of the fast transient outward K+ current IA in pyramidal neurons in (rat) primary visual cortex. J Neurosci, 2005. 25(40): p. 9185–94.

32. Kim, J., et al., Regulation of dendritic excitability by activity-dependent trafficking of the A- type K+ channel subunit Kv4.2 in hippocampal neurons. Neuron, 2007. 54(6): p. 933–47.

33. Gutman, G.A., et al., International Union of Pharmacology. LIII. Nomenclature and molecular relationships of voltage-gated potassium channels. Pharmacol Rev, 2005. 57(4): p. 473–508.

34. Laurette, P., et al., Transcription factor MITF and remodeller BRG1 define chromatin organisation at regulatory elements in melanoma cells. Elife, 2015. 4.

35. Calero-Nieto, F.J., et al., Key regulators control distinct transcriptional programmes in blood progenitor and mast cells. EMBO J, 2014. 33(11): p. 1212–26.

36. Strub, T., et al., Essential role of microphthalmia transcription factor for DNA replication, mitosis and genomic stability in melanoma. Oncogene, 2011. 30(20): p. 2319–32.

37. Shoag, J., et al., PGC-1 coactivators regulate MITF and the tanning response. Mol Cell, 2013. 49(1): p. 145–57.

38. Bepari, A.K., et al., Visualization of odor-induced neuronal activity by immediate early gene expression. Bmc Neuroscience, 2012. 13.

39. Fontanals-Cirera, B., et al., Harnessing BET Inhibitor Sensitivity Reveals AMIGO2 as a Melanoma Survival Gene. Molecular Cell, 2017. 68(4): p. 731-+.

40. Walke, W., G. Xiao, and D. Goldman, Identification and characterization of a 47 base pair activity-dependent enhancer of the rat nicotinic acetylcholine receptor delta-subunit promoter. J Neurosci, 1996. 16(11): p. 3641–51.

41. Kim, T.K., et al., Widespread transcription at neuronal activity-regulated enhancers. Nature, 2010. 465(7295): p. 182–U65.

42. Malik, A.N., et al., Genome-wide identification and characterization of functional neuronal activity-dependent enhancers. Nature Neuroscience, 2014. 17(10): p. 1330–1339.

43. Vierbuchen, T., et al., AP-1 Transcription Factors and the BAF Complex Mediate Signal-Dependent Enhancer Selection. Mol Cell, 2017. 68(6): p. 1067–1082 e12.

44. Yang, M. and J.N. Crawley, Simple behavioral assessment of mouse olfaction. Curr Protoc Neurosci, 2009. Chapter 8: p. Unit 8 24.

45. Hayar, A., et al., External tufted cells: a major excitatory element that coordinates glomerular activity. J Neurosci, 2004. 24(30): p. 6676–85.

46. Gire, D.H., et al., Mitral Cells in the Olfactory Bulb Are Mainly Excited through a Multistep Signaling Path. Journal of Neuroscience, 2012. 32(9): p. 2964–2975.

47. Fadool, D.A., et al., Kv1.3 channel gene-targeted deletion produces “Super-Smeller Mice” with altered glomeruli, interacting scaffolding proteins, and biophysics. Neuron, 2004. 41(3): p. 389–404.

48. Nunez-Parra, A., et al., Disruption of centrifugal inhibition to olfactory bulb granule cells impairs olfactory discrimination. Proceedings of the National Academy of Sciences of the United States of America, 2013. 110(36): p. 14777–14782.

49. Nunes, D. and T. Kuner, Disinhibition of olfactory bulb granule cells accelerates odour discrimination in mice. Nature Communications, 2015. 6.

50. Mombaerts, P., et al., Visualizing an olfactory sensory map. Cell, 1996. 87(4): p. 675–86.

51. Titley, H.K., N. Brunel, and C. Hansel, Toward a Neurocentric View of Learning. Neuron, 2017. 95(1): p. 19–32.

52. Pozo, K. and Y. Goda, Unraveling mechanisms of homeostatic synaptic plasticity. Neuron, 2010. 66(3): p. 337–51.

53. Hilgenberg, L.G. and M.A. Smith, Preparation of dissociated mouse cortical neuron cultures. J Vis Exp, 2007(10): p. 562.

54. Desmaisons, D., J.D. Vincent, and P.M. Lledo, Control of action potential timing by intrinsic subthreshold oscillations in olfactory bulb output neurons. J Neurosci, 1999. 19(24): p. 10727–37.

55. Fock, V., et al., Subcellular localization and stability of MITF are modulated by the bHLH-Zip domain. Pigment Cell Melanoma Res, 2018.

55. Livak, K.J. and T.D. Schmittgen, Analysis of relative gene expression data using real-time quantitative PCR and the 2(T)(-Delta Delta C) method. Methods, 2001. 25(4): p. 402–408.

57. Lehmkuhl, A.M., E.R. Dirr, and S.M. Fleming, Olfactory Assays for Mouse Models of Neurodegenerative Disease. Jove-Journal of Visualized Experiments, 2014(90).

